# Development of Physiologically Based Liver Distribution Model that Incorporates Intracellular Lipid Partitioning and Binding to Fatty Acid Binding Protein 1 (FABP1)

**DOI:** 10.64898/2026.01.17.700130

**Authors:** Yue Winnie Wen, Nina Isoherranen

## Abstract

Steady-state volume of distribution (V_ss_) can be predicted using tissue-to-plasma partition coefficients (K_p_) and tissue volumes. K_p_ values are important components of physiologically based pharmacokinetic (PBPK) models, allowing for estimation of distribution kinetics and simulation of concentration-time profiles. Many *in silico* approaches have been developed to predict tissue K_p_ values based on physicochemical processes that govern drug distribution. However, these methods frequently over- or under-predict tissue K_p_ values, highlighting the need to consider additional mechanisms that can impact drug distribution kinetics. Many drugs have been shown to bind to rat and human fatty acid binding proteins (FABPs) *in vitro* but the impact of this binding to drug distribution has not been incorporated into K_p_ predictions. We hypothesized that incorporating intracellular protein binding into tissue K_p_ predictions will improve K_p_ prediction accuracy. Using liver as a model organ, four physiologically based dynamic liver distribution models (LDMs) were developed to assess the role of distribution processes in K_p_ predictions. The developed LDMs incorporated known distribution mechanisms and intracellular drug binding to liver FABP (FABP1). The liver K_p_ values for drugs that bind to FABP1 were accurately predicted using the LDM that incorporates lipid partitioning, albumin distribution, and FABP1 binding but not using LDMs without FABP1 binding. Human FABP1 expression was quantified in 61 human livers and the interindividual variability in tissue FABP1 binding was incorporated into tissue K_p_ predictions. These simulations showed that intracellular FABP1 binding can cause interindividual variability in K_p_ values and result in concentration dependent tissue distribution.

**Significance Statement:** This study shows that incorporating intracellular protein binding such as binding to FABP1 into tissue K_p_ predictions improves accuracy of the predictions. The novel dynamic LDM can be extrapolated to other organs of interest and integrated into full-body PBPK models to predict drug distribution kinetics. With dynamic and saturable distribution mechanisms incorporated into a PBPK model, nonlinear distribution kinetics can be simulated for various drugs.

## 1. Introduction

Steady-state volume of distribution (V_ss_) is an important pharmacokinetic parameter that describes the extent of drug distribution to tissues when distribution equilibrium is achieved.^1^ V_ss_ can be measured experimentally during intravenous infusion at steady state^2–5^ or predicted using one of the mechanistic static models proposed to predict V_ss_ in humans.^6,7^ A key component in *in silico* V_ss_ prediction is consideration of tissue-to-plasma partition coefficients at distribution equilibrium (K_p_) in the context of drug-specific and physiological organ properties.^8,9^ By including organ blood flows, organ volumes, and organ specific K_p_ values into a full-body physiologically based pharmacokinetic (PBPK) model, the rate and extent of distribution to specific organs can be predicted. When combined with clearance parameters, the overall plasma concentration-time curve can be simulated even based on just *in vitro* data.^2^ Yet, methods for accurate prediction of human tissue K_p_ values are lacking for fit-for-purpose PBPK modeling and prediction of V_ss,_ drug half-life and tissue distribution kinetics.^10^

Several mechanistic static *in silico* approaches have been published to predict tissue K_p_ values and V_ss_.^3,4,11,12^ The Poulin and Theil method incorporates drug distribution to tissue water, partitioning to neutral lipids and phospholipids, and drug-albumin binding.^4^ This method was further refined by incorporating the partitioning equilibria of ionizable drugs across plasma and tissue compartments with different pH values.^3,11^ This latter method is commonly referred to as the Rodgers and Rowland method and it has been the most comprehensive and commonly used mechanistic approach for predicting tissue unbound K_p_ (K_p,u_).^3,11^ Recently, another approach, utilizing *in vitro* measured fraction unbound in liver microsomes (f_u,mic_), was proposed to better capture lipid interactions in the tissue, allowing for better predictions of lipid partitioning in various organs.^12^ Despite the overall success in predicting tissue partitioning and V_ss_ for drugs in different chemical classes, over 30% of K_p,u_ or K_p_ values for drugs remain poorly predicted based on comparison of experimentally observed and predicted values for various tissues.^3,11,12^ This highlights the need to expand and improve existing methods of K_p_ predictions.^2^

Oftentimes, static tissue K_p_ values are input alongside with tissue blood flow and size into a dynamic PBPK model, allowing for prediction of drug distribution kinetics. However, distribution mechanisms can be dynamic and some of the processes governing drug distribution can be saturable or concentration dependent. None of the existing tissue distribution prediction methods incorporate dynamic distribution mechanisms like intracellular protein binding to account for tissue distribution. Under steady-state conditions where free (unbound) unionized drug concentrations are equal on both sides of a biomembrane, intracellular protein binding will result in higher drug partitioning into the tissue and higher total tissue concentration (C_t_) than what would be expected in its absence. This in turn will increase the tissue K_p_ value.

Recent studies have shown that many drugs bind to rat and human fatty acid binding proteins (FABPs) *in vitro*.^13–15^ This is important as FABPs have broad ligand specificity and are highly expressed in different tissues across species, constituting large percentage of cytosolic protein.^16–18^ We hypothesized that intracellular drug-FABP binding will increase tissue drug concentrations and lead to a higher K_p_ values than what would be predicted from simple lipid and ion partitioning of the drug. To test this hypothesis, we examined drug distribution into the liver and different methods of K_p_ predictions for a series of drugs that bind to human liver FABP1.^15^ The aim of this study was to develop a physiologically based liver distribution model (LDM) that allows prediction of drug liver K_p_ and liver cytosolic drug concentration (C_cytosol_) in individual patients using drug specific lipid partitioning, *in vitro* FABP1 binding affinity, and FABP1 expression level quantified in human livers. An outline of the study is shown in **Figure 1**. The developed distribution model can be adapted to any other organ in the body and modified to incorporate other binding proteins and tissue targets.

**Figure 1.**
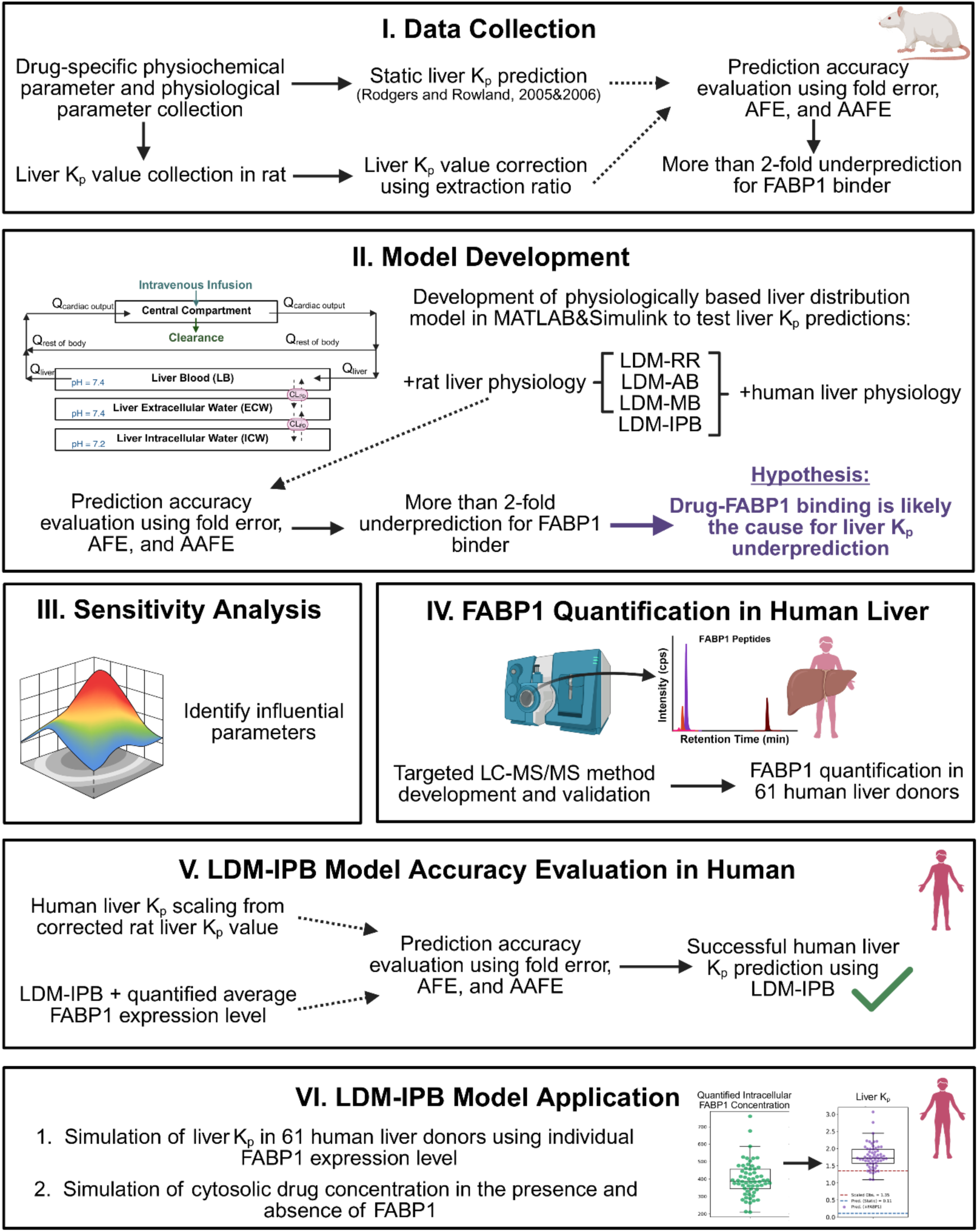
The overall workflow for developing, verifying, and applying the liver distribution model for the simulation of FABP1-binding dependent liver K_p_. FABP1, liver fatty acid binding protein; K_p_, tissue- to-plasma partition coefficient; AFE, average fold error; AAFE, absolute average fold error; LDM-RR, liver distribution model as a dynamic version of Rodgers and Rowland method; LDM-AB, liver distribution model with mechanistic albumin binding in extracellular water space; LDM-MB, liver distribution model with lipid partitioning predicted using microsomal binding data; LDM-IPB, liver distribution model with intracellular protein binding; LC-MS/MS, liquid chromatography-tandem mass spectrometry. Created in BioRender. Wen, W. (2026) https://BioRender.com/lf20yws

## 2. Material and Methods

### 2.1 Chemicals and reagents

Kanamycin, Trizma base, sodium chloride, sodium phosphate, potassium phosphate, benzonase, butanol, thrombin, Rosetta 2 *E. coli*, Coomassie Brilliant Blue R, iodoacetamide, and EDTA-free protease inhibitor cocktail were purchased from Millipore-Sigma (St. Louis, MO). Tryptone, yeast extract, isopropyl-β-d-1-thiogalactopyranoside, phenylmethyl sulfonyl fluoride, dithiothreitol, imidazole, sodium dodecyl sulfate, Pierce bicinchoninic acid (BCA) protein assay, high-performance liquid chromatography and mass spectrometry (Optima) grade acetonitrile, methanol, water and formic acid were purchased from Thermo Fisher Scientific (Waltham, MA). Lipidex- 5000 suspension was purchased from Perkin Elmer Inc (Waltham, MA, USA). Glycine and Mini-PROTEAN TGX protein gels were purchased from Bio-Rad (Hercules, CA). Mass spectrometry grade recombinant trypsin was ordered from Promega (Madison, WI). Stable isotope–labeled FABP1 peptide (SIL-peptide, F[^13^C_9_^15^N]TITAGSK, +10 Da) was ordered from Thermo Fisher Scientific (Waltham, MA) and used as internal standard for FABP1 quantification.

### 2.2 Data collection for *in vivo* distribution kinetics, K_p_ values, and tissue composition

Atenolol, (R)-propranolol, and (S)-propranolol were used as control drugs as they have weak-to-no binding to rat FABP1.^14^ Diclofenac, gemfibrozil, pioglitazone, and tolbutamide were chosen as model drugs as they are weak acids and their binding affinity to human FABP1 has been characterized.^15^ Drug specific parameters (pKa, log octanol-to-water partition coefficient (logP), blood-to-plasma ratio (BP), fraction unbound in plasma (f_u,p_) and microsomes (f_u,mic_), drug-FABP1 equilibrium dissociation constant (K_d,FABP1_), and drug-albumin equilibrium dissociation constant (K_d,albumin_)), and observed human and rat peak plasma concentrations (C_max_) were collected from public databases (e.g., DrugBank, PubChem) and literature.^19–23^ When available, experimentally measured values were used instead of *in silico* predictions. These values are summarized and listed in **Table 1**.

**Table 1.** Drug specific input parameters for prediction of liver K_p_.

| Drug<br>(references) | Atenolol<br>(19,49–53) | (R)-<br>Propranolol<br>(11,19,30,54) | (S)-<br>Propranolol<br>(11,19,30,54) | Diclofenac<br>(12,15,19,20,55,56) | Gemfibrozil<br>(15,19,21,57) | Pioglitazone<br>(15,19,22,58,59) | Tolbutamide<br>(15,19,23,56,60,61) |
| --- | --- | --- | --- | --- | --- | --- | --- |
| Human $f_{u,p}$ | 0.95 | 0.13 | 0.18 | 0.005 | 0.03 | 0.01 | 0.065 |
| Rat $f_{u,p}$ | 0.97 | 0.017 | 0.13 | 0.009 | 0.03<sup>a</sup> | 0.01 | 0.022 |
| logP | 0.38 | 3.65 | 3.65 | 4.26 | 4.39 | 3.33 | 2.34 |
| BP | 1.11 | 0.77 | 1.29 | 0.55 | 0.55<sup>b</sup> | 0.55<sup>b</sup> | 0.6 |
| pKa | 9.55<sup>c</sup> | 9.5<sup>c</sup> | 9.5<sup>c</sup> | 3.99<sup>d</sup> | 4.42<sup>d</sup> | 5.19<sup>d</sup> | 5.27<sup>d</sup> |
| $f_{u,mic}$<sup>e</sup> | 0.99 | 0.32 | 0.27 | 1 | 0.52 | 1 | 0.99 |
| $K_{d,FABP1}$<br>(μM) | 717 | No binding | No binding | 2.5 | 3.6 | 1 | 20 |
| $K_{d,albumin}$<br>(μM) | <sup>f</sup> | <sup>f</sup> | <sup>f</sup> | 2.0 (site I);<br>16.7 (site II) | 9.34<br>(site I & II) | 2.94 (site I);<br>5.88 (site II) | 21<br>(site I, II, &<br>III) |
| Human $C_{max}$<br>(μM) | 1.05 | 0.19 | 0.19 | 4.1 | 80 | 3.2 | 196 |
| Rat $C_{max}$<br>(μM) | 1.1 | 0.19<sup>g</sup> | 0.19<sup>g</sup> | 169 | 10 | 3 | 766 |

<sup>a</sup>The $f_{u,p}$ from human was used as no experimental data in rat was available.
<sup>b</sup>No experimental BP values are available for gemfibrozil and pioglitazone. A BP of 0.55 (1-hematocrit) was assumed due to their physicochemical properties.
<sup>c</sup>pKa for bases.
<sup>d</sup>pKa for acids.
<sup>e</sup>Fraction unbound in microsomes was corrected to 1 mg/mL using previously described equation<sup>12,62</sup> and is reported in the table as $f_{u,mic}$ for 1 mg/mL microsomal concentration.
<sup>f</sup>Due to minimal reported albumin binding for the basic drugs analyzed, $K_{d,albumin}$ was not collected.
<sup>g</sup>Due to lack of reported maximum concentration in the reported rat studies, same $C_{max}$ as observed in human was assumed.<sup>11,19</sup>

To collect information on liver distribution kinetics for atenolol, (R)- and (S)-propranolol, diclofenac, gemfibrozil, pioglitazone, and tolbutamide, literature published by June 2025 was searched in PubMed and Google Scholar databases with key terms of “drug name”, “intravenous infusion”, “pharmacokinetic study”, “liver distribution”, and “liver concentration”. No reports of human liver K_p_ values for above drugs were identified. All seven drugs had liver distribution data available in rat (**Table S1-1**). Hence, observed liver K_p_ values in rat were collected. Only studies that described single drug dosing in healthy animals and provided drug dosing information and tissue and plasma concentrations were included. If data were only shown in graphical format, numerical values were extracted using WebPlotDigitizer (version 4.6). If steady state IV infusion data was available, it was used for K_p_ values. If infusion data was not available, the tissue-to-plasma AUC ratio was used for K_p_ calculation. If this could not be done, individual time point liver-to-plasma concentration ratios were calculated and used as K_p_ values. No correction for terminal phase distribution was implemented due to the low clearance (**Table S1-2)** and limited distribution of the model drugs.

Liver K_p_ values at steady state after IV infusion were available for (R)- and (S)-propranolol (reported after correction for hepatic extraction) and for tolbutamide. For atenolol, liver and plasma AUC_0-inf_ were reported. For diclofenac, liver and plasma concentration versus time data were available and AUC_0-inf_ was estimated using non-compartmental analysis (NCA) in Phoenix (V8.4.0). For atenolol and diclofenac liver K_p_ was calculated from these data as the liver-to-plasma AUC_0-inf_ ratio. For gemfibrozil and pioglitazone full AUC_0-inf_ data could not be estimated. Instead, an apparent K_p_ was calculated as the average of measured liver-to-plasma concentration ratios reported at multiple time points **(Table S1-1)**.

For liver K_p_ predictions, rat and human liver physiological parameters were extracted from literature.^3,4,24–29^ The values included fractional tissue volumes of extracellular water space (f_ECW_), intracellular water space (f_ICW_), neutral lipid (f_NL_) and neutral phospholipid (f_NP_); fractional plasma volume of neutral lipid (f_p,NL_) and fractional plasma volume of neutral phospholipid (f_p,NP_), pH in plasma (pH_p_), pH of intracellular water space (pH_ICW_), and albumin concentration in liver and plasma (C_albumin,liver_, C_albumin,p_). The specific values are summarized in **Table S1-3**. In addition to physiological parameters detailed in **Table S1-3**, red blood cell pH of 7.22, liver acidic phospholipids of 4.56 mg/g, fractional blood cell volume of neutral lipid of 0.0017, fractional blood cell volume of neutral phospholipid of 0.0029, and fractional blood cell volume of intracellular water of 0.603 were used as reported^30^ for prediction of rat liver K_p_ for the basic drugs atenolol, and (R)-, and (S)-propranolol.

### 2.3 Correction of observed rat liver K_p_ values for metabolism in the liver and scaling of K_p_ values to human liver

Since liver is an eliminating organ and sampling in the rat studies was not from local liver blood, the observed liver K_p_ was corrected for liver clearance (K_p,corrected,rat_) as previously described^11,30,31^ using **equation 1**:

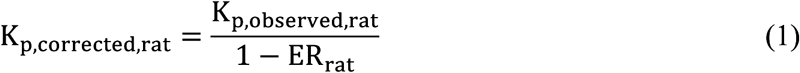

and assuming the well-stirred model of the liver. This is a reasonable assumption as none of the drugs evaluated had a high extraction ratio. The literature reported rat hepatic extraction ratio (ER_rat_) (**Table S1-2)** of the model drugs was used.

As no observed human liver K_p_ values were available, the observed K_p,corrected,rat_ values (**Table 2**) were scaled to human liver K_p_ (K_p,scaled,human_) using **equation 2** as previously described^6^ and assuming liver tissue unbound fractions are the same in human and rat.

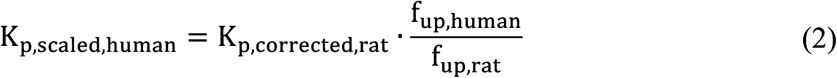

**Table 2.** Summary of the rat liver K_p_ values observed and predicted using static models or dynamic liver distribution models (LDMs) constructed based on known distribution mechanisms. The corrected liver K_p_ data in rats were estimated from *in vivo* studies and adjusted for hepatic extraction ratios. The three LDMs were the Rodgers and Rowland model (LDM-RR), the mechanistic albumin binding model (LDM-AB) and the microsomal binding model (LDM-MB).

| Drug | | Corrected<br>Liver $K_p$<sup>11,20-</sup><br><sup>23,52</sup> | Predicted Liver $K_p$ | | | |
| --- | --- | --- | --- | --- | --- | --- |
|  |  |  | Static<br>Equation<sup>3,30</sup> | LDM-RR | LDM-AB<sup>b</sup> | LDM-MB |
| Weak-to-no<br>binding to<br>FABP1 | Atenolol | 3.0 | 4.4 | 4.4 | na | 1.6 |
|  | (R)-Propranolol | 5.6<sup>a</sup> | 4.5 | 4.5 | na | 2.4 |
|  | (S)-Propranolol | 30<sup>a</sup> | 15 | 15 | na | 24 |
|  | AFE | - | 0.84 | 0.84 | - | 0.57 |
|  | AAFE | - | 1.55 | 1.55 | - | 1.76 |
| Moderate-to-tight binding<br>to FABP1 | Diclofenac | 1.4 | 0.09 | 0.09 | 0.14 | 0.14 |
|  | Gemfibrozil | 3.6 | 0.12 | 0.12 | 0.25 | 0.97 |
|  | Pioglitazone | 1.4 | 0.09 | 0.09 | 0.19 | 0.18 |
|  | Tolbutamide | 0.30 | 0.10 | 0.10 | 0.09 | 0.10 |
|  | AFE | - | 0.08 | 0.08 | 0.13 | 0.18 |
|  | AAFE | - | 12.11 | 12.11 | 7.69 | 5.41 |

<sup>a</sup>Liver $K_p$ was reported after correction for hepatic extraction in the original reference.<sup>11</sup>
<sup>b</sup>Atenolol and (R)-, and (S)-propranolol are basic drugs that have negligible extracellular albumin binding. Hence liver $K_p$ was not simulated using LDM-AB.

### 2.4 Static rat liver K_p_ predictions

Rat liver K_p_ values were predicted for the control drugs and for the four acidic model drugs as published for bases^30^ and for weak acids^3^ using **equation 3**:

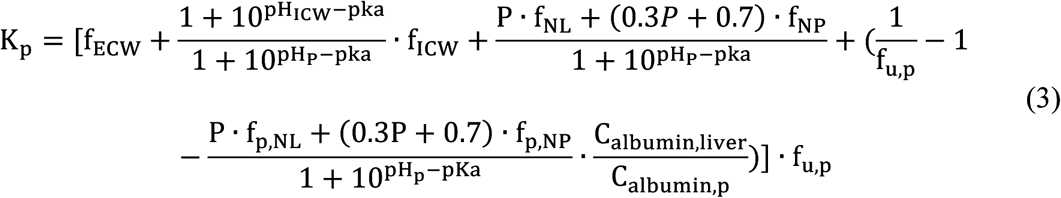

The parameters in the equation are as described above. The values listed in **Table 1 & Table S1- 3** for rat were used.

### 2.5 Development of physiologically based liver distribution models (LDM)

A 4-compartment rat physiologically based LDM (**Figure 2**) was developed in MATLAB and Simulink platform (R2025b)^32^. The model includes a circulation component and a physiologically based mechanistic LDM consisting of liver blood, extracellular water, and liver tissue. The mechanistic LDM structure is based on known rat and human physiology. Three different models (LDM-RR, LDM-AB and LDM-MB) were constructed to simulate steady-state rat liver K_p_ (**Figure 2**). The first one (LDM-RR) was built as a dynamic version of the original Rodgers and Rowland (RR) method^3,30^. The second model (LDM-AB) incorporated published affinity (K_d,albumin_) for drug-albumin binding (AB) to predict extracellular drug protein binding and drug concentrations in extracellular water for acidic drugs. The third model (LDM-MB) has the same structure as LDM-AB but incorporates microsomal binding (MB) data^12^ to predict lipid partitioning for all model drugs. The developed rat and human LDMs are available at GitHub (https://github.com/Isoherranen-Lab/Liver-Distribution-Model).

**Figure 2.**
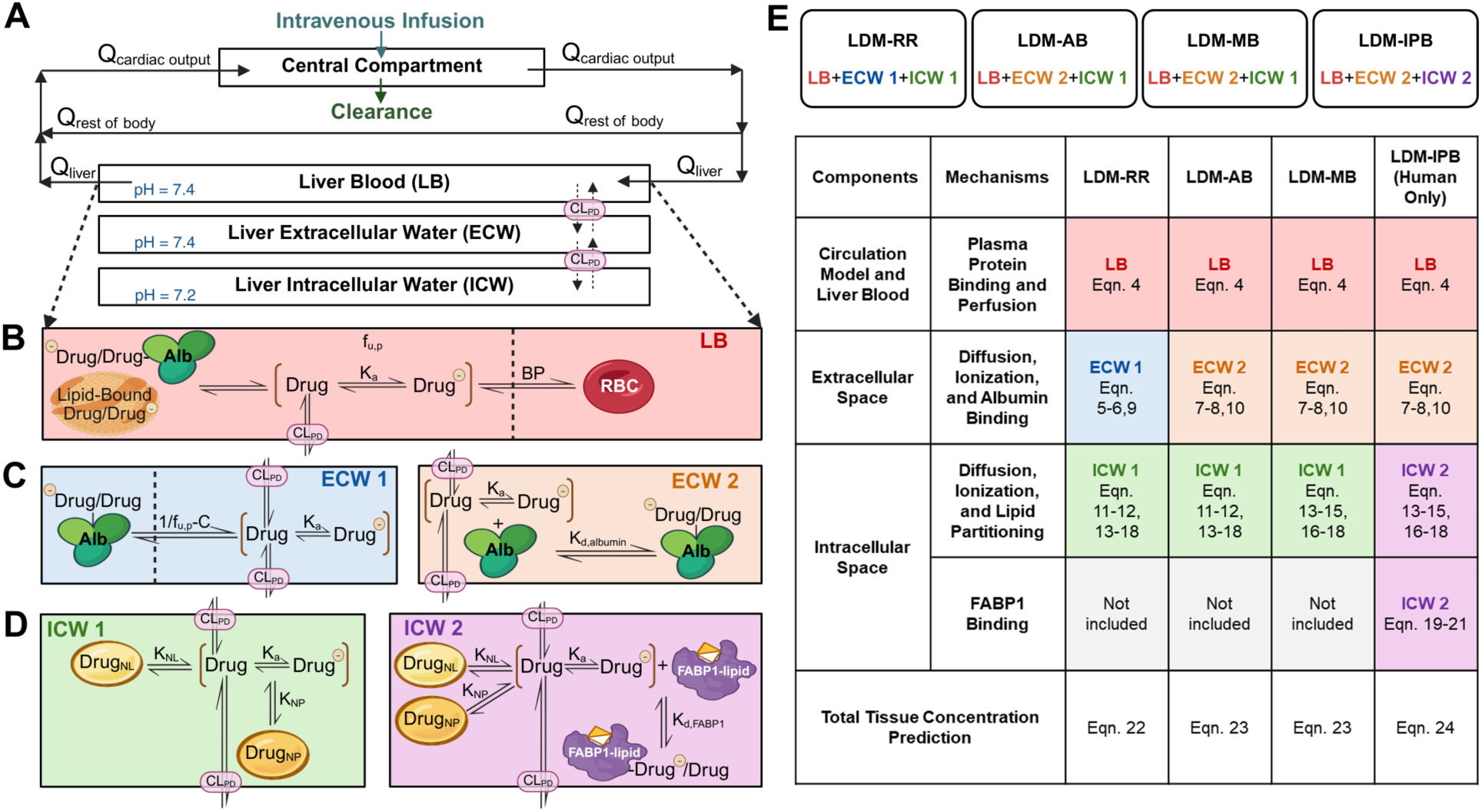
Schematic diagram of the liver distribution model (LDM). Panel A shows the overall 4- compartment model structure. Panels B-D illustrate liver blood (LB), different extracellular water (ECW) and intracellular water (ICW) models. Panel E is a summary for liver model structures of the four LDMs developed in this study with corresponding equation numbers that describe the differential equations for the model. Q_cardiac output_, blood flow to the central compartment; Q_rest of the body_, blood flow to the rest of the body; Q_liver_, blood flow to the liver; CL_PD_, passive diffusion clearance; K_a_, acid dissociation constant; f_u,p_, unbound fraction of drug in the plasma; C, a constant derived from equation 5 for drug-albumin partitioning 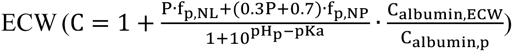, dissociation constant between albumin and drug; K_NL_, partition constant for neutral lipid binding; Drug_NL_, drug partitioned into the neutral lipid; K_NP_, partition constant for neutral phospholipid binding; Drug_NP_, drug partitioned into the neutral phospholipid. K_d, FABP1_, dissociation constant between FABP1 and drug. Created in BioRender. Wen, W. (2026) https://BioRender.com/neske0m

A simple circulation compartment was used to connect the liver blood flow out of the liver blood compartment to the liver blood flow into the liver blood compartment (**Figure 2**). Within the circulation compartment the drug may partition into red blood cells and bind to plasma proteins. Plasma lipid and protein binding, ionization of the drug, extracellular water protein binding, and lipid partitioning were included in corresponding liver compartments according to previously published mechanisms.^3,12^ Only unbound unionized drug was allowed to diffuse between liver compartments. The drug was administered as IV infusion with clearance (CL) included in the central circulation component to allow simulations of equilibrium conditions. The infusion rate (R_0_) and CL were set such that the ratio R_0_/CL recapitulated a steady-state concentration (C_ss_) corresponding to half of the reported peak concentration (C_max_) observed in the rat listed in **Table 1**.

#### 2.5.1 Liver blood compartment and drug concentrations in liver blood (LB)

The same LB compartment was used for all models developed. **Equation 4** describes the change of blood drug concentration in the circulation compartment with respect to time:

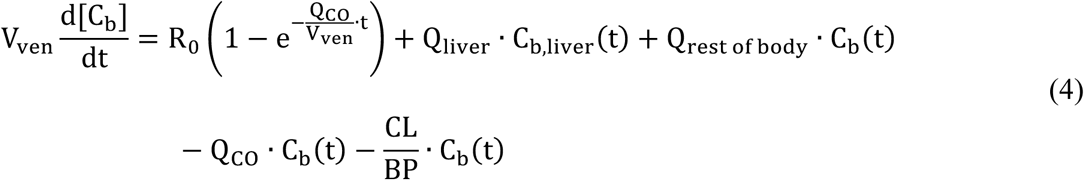

where C_b_ is venous blood drug concentration (µmol/L), V_ven_ is venous blood volume (0.0123 L)^24^, R_0_ is IV infusion rate (mg/hr); Q_liver_ is liver blood flow (0.825 L/hr)^33^, C_b,liver_ is liver blood concentration (µmol/L), Q_rest of body_ is blood flow to the rest of the body calculated as the difference between liver blood flow and cardiac output (4.575 L/hr)^24^, Q_CO_ is cardiac output (5.4 L/hr)^24^, CL is systemic plasma clearance (L/hr), and BP is blood-to-plasma ratio.

#### 2.5.2 Liver extracellular water (ECW) models

Two different liver ECW models were built to predict drug binding to albumin in ECW (**Figure 2C**). In the first model (ECW 1) the ECW partitioning of albumin-bound drug (K_albumin_) was predicted based on plasma unbound fraction using **equation 5** as previously described for weakly acidic drugs^3^:

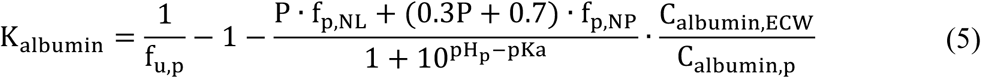

In **equation 5**, P is octanol-to-water partition coefficient, f_p,NL_ is fraction of plasma wet weight as neutral lipid (0.00149)^34^, f_p,NP_ is fraction of tissue wet weight as neutral phospholipid (0.00083)^34^, 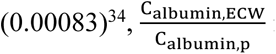 is liver extracellular water-to-plasma albumin concentration ratio (0.54)^3^, pH_p_ is the plasma pH (7.4)^27^, and pKa is the drug pK_a_. The K_albumin_ can be used in **equation 6** to describe the change of drug-albumin concentration (C_drug–albumin,ECW_ ; µmol/L) in the ECW with respect to time:

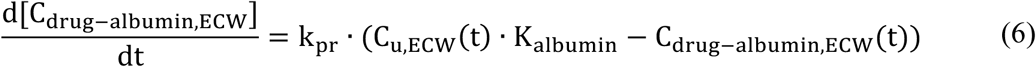

where k is the rate constant for partitioning, assumed to be 1 hr^-1^ for model drugs.

In the second model (ECW 2), experimentally measured drug-albumin binding affinity (K_d,albumin_) and C_albumin,ECW_ were used to simulate drug binding and distribution into the ECW. As drug binding to multiple binding sites on human serum albumin was reported for all model drugs (**Table 1**), multiple binding sites of albumin were included in the model. Both unionized and ionized drug were assumed to have the same K_d,albumin_. At any given time, the sum of free albumin concentration and the drug-bound albumin concentration for a given binding site equals the C_albumin,ECW_ (221 µmol/L)^3^.

Assuming drug-albumin association is diffusion rate limited^35^, the dissociation rate constant of drug from the binding site i (k_off,albumin,site i_) can be calculated from experimentally measured ^K^d,albumin,site i^:^

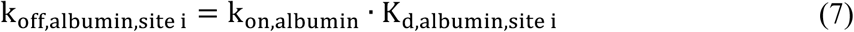

where k_on,albumin_ is the association rate constant (360,000 µM^-1^hr^-1^)^35^.

The change in C_drug–albumin,ECW_ can be expressed with unbound concentration of albumin binding site i (C_albumin,u,ECW,site i_; µmol/L) and the drug-bound concentration of albumin binding site i (Cdrug–albumin,ECW,site n; µmol/L):

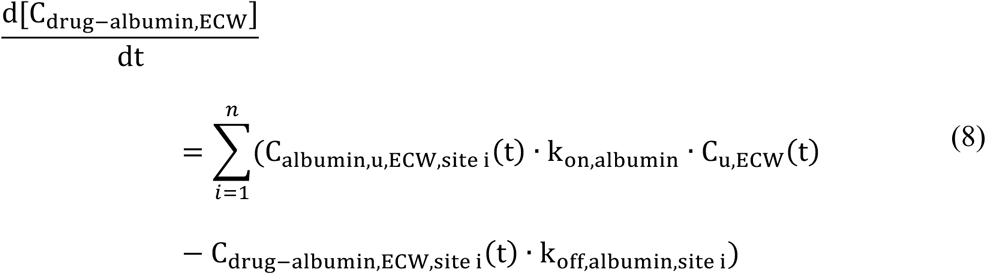

Overall, the change in C_u,ECW_ can be expressed for the two ECW models as follows:

ECW 1:

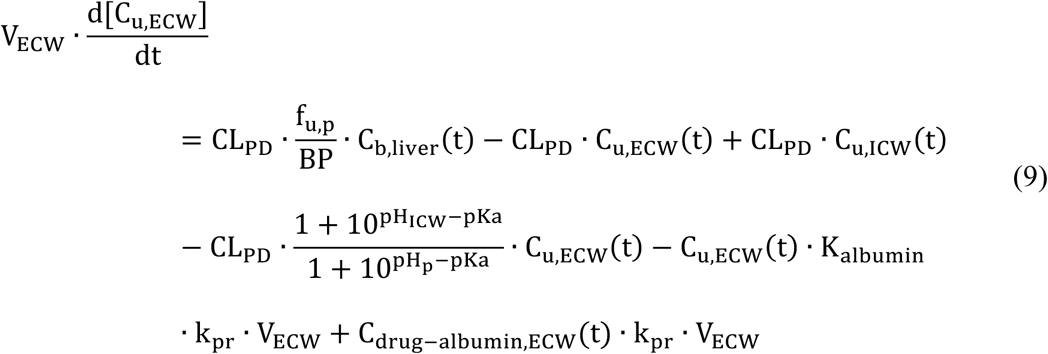

where V_ECW_ is the volume of liver ECW (0.00115 L),^24,34^ CL_PD_ is the passive diffusion clearance (a value of 100,000 L/hr greatly exceeding liver blood flow was used to ensure perfusion rate limited distribution), C_u,ICW_ is the unbound concentration in intracellular water in liver (µmol/L), and pH_ICW_ is the pH of the intracellular water (7.2).^27^

ECW2:

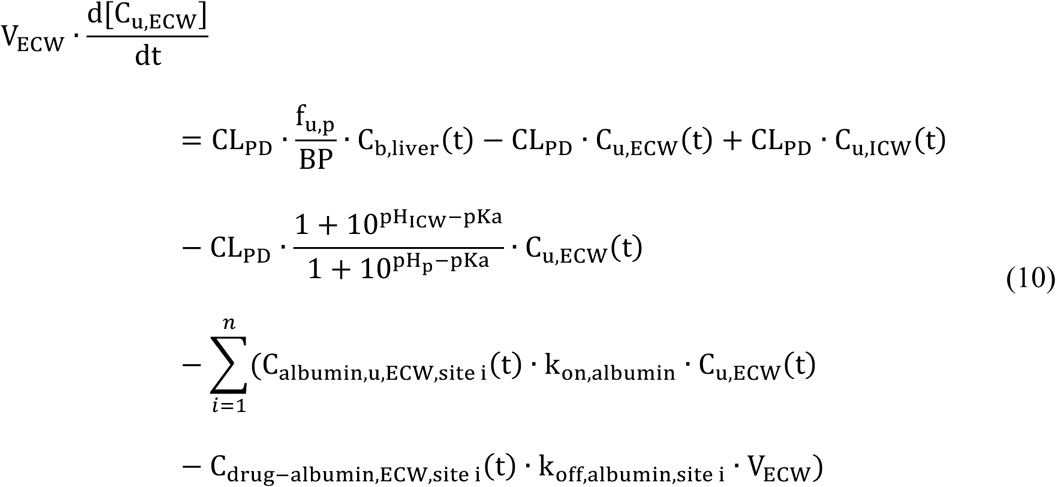

#### 2.5.3 Liver intracellular water (ICW) model without intracellular protein binding

To predict drug distribution into the liver ICW, a model structure ICW1 was built. This model incorporated two different methods for predicting lipid partitioning (LP), model LP 1 and 2 (**Figure 2D**). For weakly acidic drugs, drug partitioning into the neutral lipids (NL) and neutral phospholipids (NP) was considered and modeled relying on partition constant for NL (K_NL_) and partition constant for NP (K_NP_). K_NL_ and K_NP_ were derived based on different published mechanisms^3,12^ and used as such in this dynamic model. In LP 1, K_NL_ and K_NP_ were predicted using physiochemical properties of the drug while in LP 2 *in vitro* f_u,mic_ (**Table 1**) was used as a surrogate for natural membrane partitioning. The two methods of lipid partitioning are described in **equations 11-15**.

LP 1:

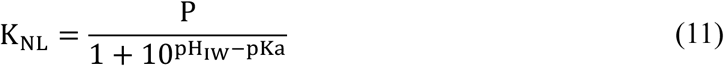

LP 2:

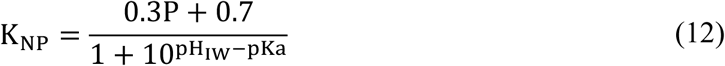

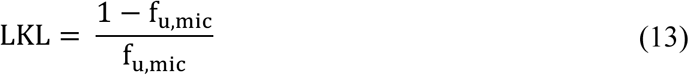

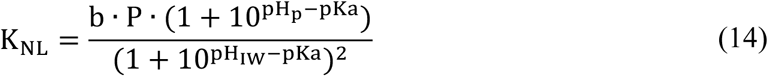

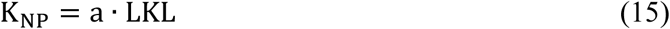

In **equations 14 and 1**5, a and b are the previously optimized constants (a = 1383; b = 0.096).^12^

For both LP 1 and LP 2, the change in drug concentration in NL (C_NL_) or NP (C_NP_) can be expressed as follows:

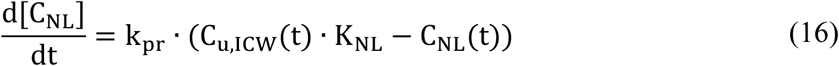

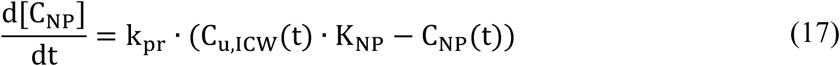

Overall, the change in C_u,ICW_ in the liver with respect to time can be described using **equation 18**:

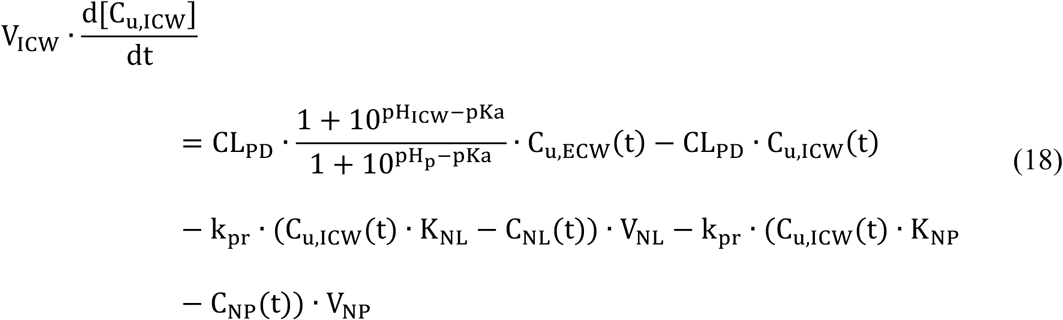

where V_NL_ is the volume of liver NL (0.0000998 L)^24,34^ and V_NP_ is the volume of liver NP (0.000219 L)^24,34^.

#### 2.5.4 Liver intracellular water (ICW) model with intracellular FABP1 binding

To predict liver drug distribution in humans, liver distribution model with intracellular protein binding (LDM-IPB) was constructed by adapting the LDM-MB with all human physiological system parameters (**Table S1-3).** For intracellular lipid partitioning ICW2 was used. For LDM- IPB, the drug-FABP1 binding was added in the ICW of the liver in addition to lipid partitioning for predicting drug liver K_p_ at steady state (**Figure 2**). At any given time, the sum of the concentrations of unbound FABP1 (C_FABP1,u_; µmol/L) and drug-bound FABP1 (C_drug–FABP1_; µmol/L) equals the total cytosolic FABP1 concentration quantified in this study (C_FABP1_; µmol/L).

Assuming the drug-FABP1 binding process is diffusion rate limited,^35^ the drug-FABP1 k_off_ (µM^-1^hr^-1^) can be expressed from K_d,FABP1_ (µmol/L):

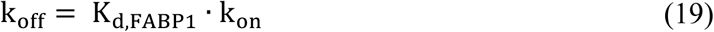

where k_on_ is the association rate constant (360,000 µM^-1^hr^-1^).^35^

Hence, the change in C_drug–FABP1_ with respect to time is described by **equation 2**0:

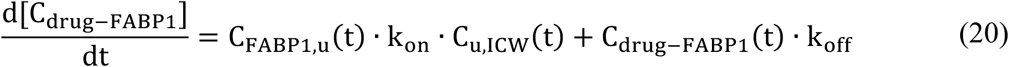

With the incorporation of FABP1 binding, the unbound ICW drug concentration can be described as:

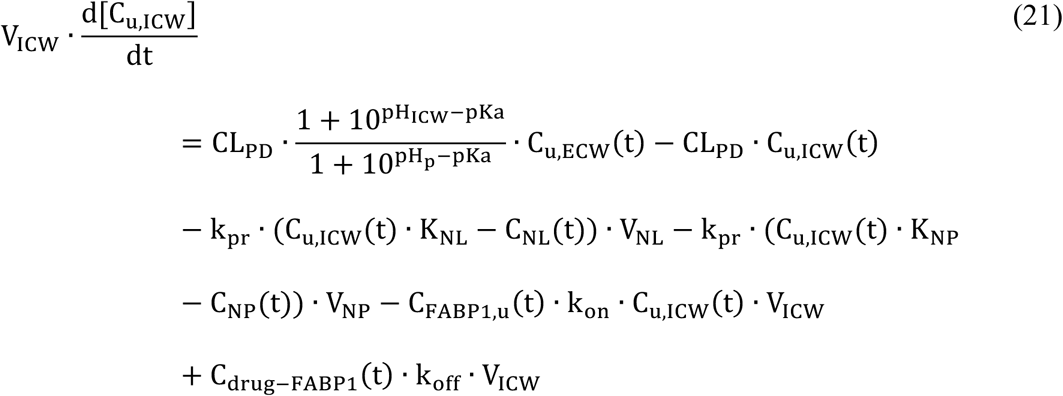

#### 2.5.5 Overall tissue concentration and liver K_p_ prediction

The liver K_p_ was calculated as the ratio of model output total drug tissue concentration (C_liver_; µmol/L) and plasma concentration (C_p_; µmol/L). Predicted C_liver_ was calculated in the different models as follows:

LDM-RR:

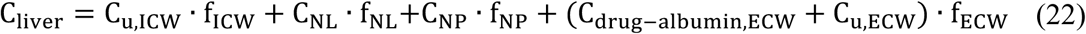

LDM-AB and LDM-MB:

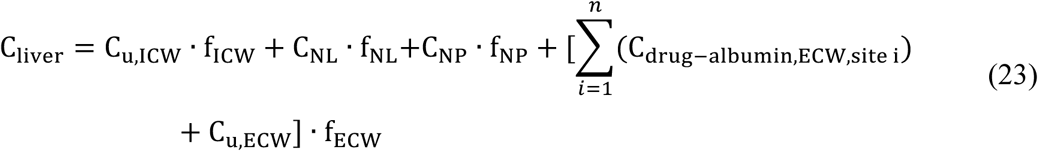

LDM-IPB:

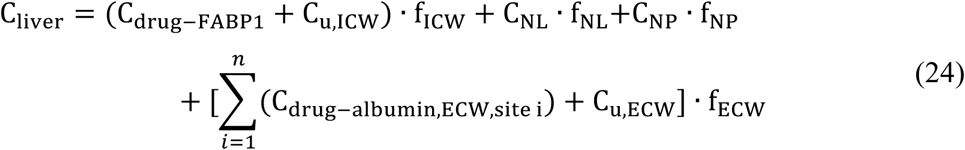

### 2.6 Assessment of prediction accuracy

To assess the prediction accuracy of the different distribution models in rat and human, the liver K_p_ values of atenolol, (R)- and (S)-propranolol or diclofenac, gemfibrozil, pioglitazone, and tolbutamide predicted at C_ss_ set as half of C_max_ were compared to K_p,corrected,rat_ or K_p,scaled,human_. The model performance was evaluated using fold error, average fold error (AFE) and absolute average fold error (AAFE) (**Eq.** 25, 26, & 27). A predetermined two-fold acceptance criterion was applied to determine whether liver K_p_ was successfully simulated against K_p,corrected,rat_ or K_p,scaled,human_.

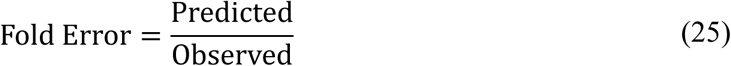

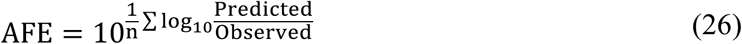

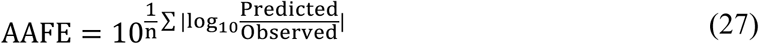

### 2.7 Sensitivity analysis for the physiologically based human LDM-MB and LDM-IPB

A sensitivity analysis was performed to identify key model input parameters influencing the predicted human liver K_p_ and C_cytosol_ in LDM-MB and LDM-IPB. A hypothetical weakly acidic drug with f_u,p_ of 0.01, logP of 3.84, BP of 0.55, pKa of 5.14, f_u,mic_ of 0.88 at 1 mg/mL human liver microsomal protein concentration, K_d,albumin_ of 2 µM with single binding site, and C_ss_ of 40 µM was used for LDM-MB assessment. In addition, intracellular FABP1 expression of 399 µM and K_d,FABP1_ of 3.6 µM were included for LDM-IPB. Each parameter of interest was varied individually over a physiochemically or physiologically plausible range while all other parameters were held constant. The covariates (range of values indicated in brackets) tested for evaluating liver K_p_ and C_cytosol_ included logP (1-5), f_u,mic_ (0.1-1), pKa (2-7), f_u,p_ (0.0001-0.05), K_d,albumin_ (2-20 µM), and C_ss_ (0.1-200 µM) for LDM-MB as well as intracellular FABP1 expression (100-900 µM) and K_d,FABP1_ (1-20 µM) for LDM-IPB.

### 2.8 Quantification of FABP1 in human liver S9 fractions (HLS9) using LC-MS/MS and scaling to human liver concentrations

Human liver samples from 61 individual donors were obtained from the University of Washington human liver bank. FABP1 expression in HLS9 from 61 livers was measured in singlet on three separate days using surrogate peptide-based LC-MS/MS protein quantification. The average of the three independent quantifications is reported. HLS9 protein yield per gram of liver tissue was calculated for each donor from protein concentrations quantified in HLS9 using BCA assays. The measured FABP1 concentration as nmol FABP1 per milligram of S9 protein was extrapolated to liver expression using the HLS9 yield. Assuming a tissue density of 1 mL/g, intracellular FABP1 concentration (µM) was estimated using tissue FABP1 expression level and f_ICW_of 0.573 (**Table S3-1**).

For FABP1 quantification, the unique signature peptide FTITAGSK (**Table S3-2**) was measured using a Sciex 5500 QTrap linear ion trap mass spectrometer (AB Sciex, Foster City, CA) coupled to an Agilent 1290 LC (Agilent, Santa Clara, CA). Details regarding expression and purification of the FABP1 reference protein (**Method S3-1**), preparation of mouse liver S9 (MLS9) and HLS9 fractions(**Method S3-2**), selection of surrogate peptide (**Method S3-3, Figure S3-1&S3-2, Table S3-2**), optimization of digestion conditions (**Method S3-4, Figure S3-3&S3-4**), and evaluation of matrix effects are described (**Method S3-4, Figure S3-5**). Heavy labeled internal standard peptide (F[^13^C_9_^15^N]TITAGSK) was ordered from Thermo Fisher Scientific (± 5–10% accuracy and >98% purity).

Peptides were separated using an Aeris peptide XB-C18 column (50 × 2.1 mm, 1.7 mm particle size), and a SecurityGuard Ultra LC C18-peptide cartridge (Phenomenex, Torrance, CA). Gradient elution at 400 μL/min with H_2_O (A) and acetonitrile (B) both with 0.1% formic acid was used at 40°C as follows: 3% B until 3.5 min, increased to 40% B by 12.0 min, then to 95% B by 12.1 min and kept at 95% B until 14.0 min, returning to 3% B at 14.1 min with run time of 17 min. The peak areas were integrated using Skyline-daily^36,37^. For quantification, the ratio of FTITAGSK peptide peak area to peptide area of 20 nM F[^13^C_9_^15^N]TITAGSK was used to construct standard curves and calculate FABP1 concentrations in individual HLS9.

Protein digestion was done as previously described.^38^ Calibration curve or quality control (QC) samples for FABP1 quantification were prepared by spiking purified human FABP1 into 0.2 mg/mL MLS9 (final concentration 50–1000 nM for standard curve and 50, 120, 320, 540 nM for lower limit of quantitation (LLOQ), low QC, medium QC, and high QC). To 40 µL of 0.2 mg/mL HLS9 and MLS9, 20 nM F[^13^C_9_^15^N]TITAGSK internal standard was added and the samples were incubated with 8 µL of 100 mM dithiothreitol followed by addition of 20 µL of 100 mM ammonium bicarbonate (pH 7.8) and incubation for 20 minutes at room temperature. Then, 10 µL of 10% sodium deoxycholate in 100 mM ammonium bicarbonate (pH 7.8) were added. Proteins were denatured at 95°C for 5 minutes in a ThermoMixer (Thermo Fisher Scientific, Waltham, MA). 8 µL of 200 mM iodoacetamide were added to alkylate cysteine residues. The samples were incubated at room temperature for 20 minutes in the dark. 4 µL of 0.2 µg/µL trypsin were then added to digest the protein at 37 °C for 5 hours at a 1:10 trypsin/protein (w/w) ratio. Trypsin digestion was stopped by the addition of 40 µL ice-cold ACN containing 8% trifluoroacetic acid. Samples were centrifuged at 3,400 g at 4 °C for 60 minutes. The supernatant was transferred to a new 96-well plate for LC-MS/MS analysis as described above.

The assay was validated according to the FDA guidance^39^, including linearity, LLOQ, precision, accuracy, stability of digested surrogate peptides, and carryover. Signal linearity was first confirmed by serial dilution of digested purified protein, equivalent to starting at 300 nM of purified FABP1. Instrument variance was assessed by calculating coefficient of variance (CV) from six replicate injections.

Five replicates per QC levels were digested per day for four independent experiments and analyzed across ten days to determine inter-day variance. Intra-day variance was calculated from five QC samples per concentration within a day and the mean intra-day variance from ten days is reported. Aliquots of a pool of HLS9 from 61 donors were prepared and run in five replicates with every run as pooled QCs. Freeze-thaw and autosampler stability of digested peptides was determined by quantifying QC samples that were either frozen at – 20°C for two freeze-thaw cycles or kept at 10°C in autosampler for 24-hr and 40-hr, respectively (**Table S3-3**).

### 2.9 Prediction of individual human liver K_p_ values

The liver distribution of diclofenac, gemfibrozil, pioglitazone, and tolbutamide was predicted using the LDM-IPB, previously reported K_d,FABP1_ (**Table 1**) for the four drugs^15^ and the average human liver FABP1 expression level quantified in this study. The simulated human liver K_p_ values were compared to K_p,scaled,human_.

Upon acceptance of fold error, AFE, and AAFE for mean distribution values, a population-based simulation approach was employed. The individual donor intracellular FABP1 concentrations quantified using the LC-MS/MS method were incorporated into the model to simulate liver K_p_ for the 61 donors. The interindividual variability in liver K_p_ due to varying FABP1 expression level was predicted for all the selected drugs.

### 2.10 Simulation of C_cytosol_ with and without FABP1 binding

To examine the impact of drug-FABP1 binding on C_cytosol_, simulations were performed for the 4 model drugs using the LDM-IPB and using **equation 28** adapted from the published static equation^3^ (**Eq.3**).

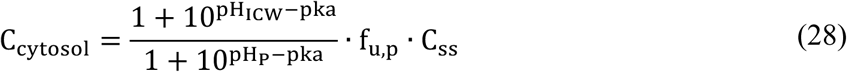

For the LDM-IPB, C_cytosol_ is reported as the sum of the free drug concentration and the concentration of drug-FABP1 complex in ICW. A quantified population average FABP1 expression level from 61 individual donors was used for the simulations with drug specific K_d,FABP1_ considered.

## 3. RESULTS

### 3.1 Prediction of rat liver K_p_

The rat liver K_p_ values for atenolol, (R)-propranolol, (S)-propranolol, diclofenac, gemfibrozil, pioglitazone, and tolbutamide were estimated from literature data and corrected for the impact of liver metabolism (**Table 2**). The rat liver K_p_ values predicted using the static equations^3,30^ were within 2-fold of the observed for drugs with weak-to-no binding to FABP1, but underpredicted by 3- to 30-fold for the acidic drugs with moderate-to-tight binding to FABP1 (**Table 2**). To explore the reasons for the underprediction, a dynamic 4-compartment LDM was developed based on known physiology (**Figure 2**). The dynamic model using the same distribution concepts as the previous static model (LDM-RR) resulted in identical K_p_ prediction values as the static equation (**Table 2 & Figure 3**) confirming model integrity but suggesting a systematic underprediction of K_p_ values for acidic drugs.

**Figure 3.**
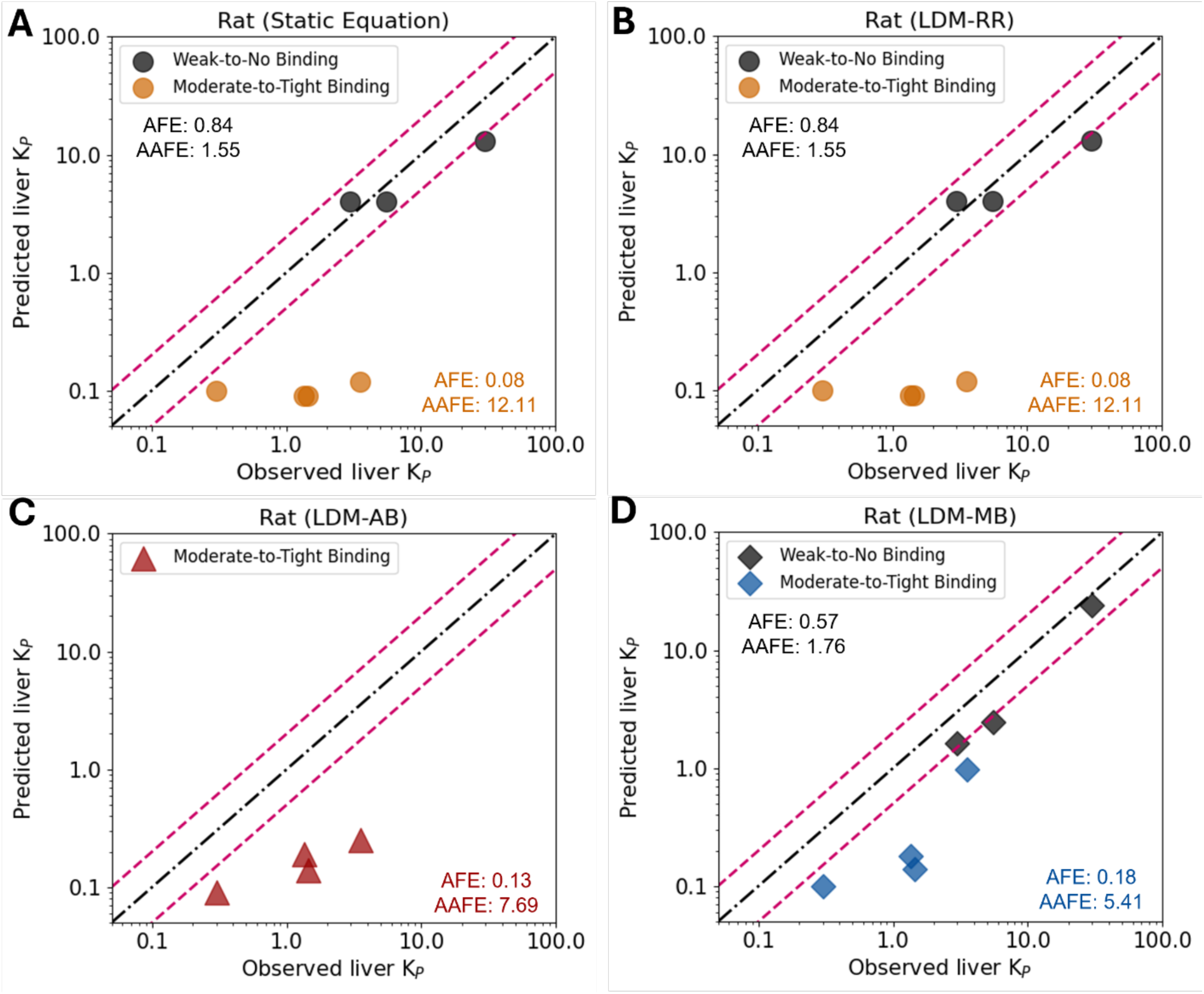
Observed versus predicted rat liver K_p_ values for drugs with weak-to-no binding to FABP1 (atenolol, (R)-propranolol, and (S)-propranolol, black symbols) and moderate-to-tight binding to FABP1 (diclofenac, gemfibrozil, pioglitazone, and tolbutamide, orange, red, and blue symbols). Panel A shows the observed versus predicted rat liver K_p_ using static equation for bases^30^ or acids^3^. Panel B shows the observed versus predicted rat liver K_p_ using liver distribution model with Rodgers and Rowland mechanisms^3,30^ (LDM-RR). Panel C shows the observed versus predicted rat liver K_p_ using liver distribution model with mechanistic albumin binding (LDM-AB). Panel D shows the observed versus predicted rat liver K_p_ using liver distribution model with microsomal binding (LDM-MB). The black dashed line is the line of unity when the observed and predicted values are the same. The pink dash lines represent two-fold error. AFE is the average fold error and AAFE is the absolute average fold error.

To explore the reasons for the underprediction, the distribution characteristics of diclofenac, gemfibrozil, pioglitazone, and tolbutamide were further evaluated. All 4 model drugs are highly bound to albumin in plasma.^40^ To test whether poor estimation of drug-albumin binding kinetics and albumin distribution to the liver could explain the underprediction of liver K_p_, a refined ECW albumin binding model (LDM-AB) was developed (**Figure 2**). Rat liver K_p_ values were then predicted using experimentally reported K_d,albumin_ (**Table 1**). This resulted in improved prediction accuracy (**Figure 3)**, but the observed rat liver K_p_ values were still 3- to 14-fold higher than the predicted K_p_ values (**Table 2**). The AFE and AAFE for the rat liver K_p_ prediction using the LDM- AB of the 4 drugs were 0.13 and 7.69 respectively (**Table 2 & Figure 3**).

We hypothesized that the liver K_p_ prediction error may be due to underprediction of lipid partitioning for the test drugs as the prediction of lipid partitioning in the model is based on physicochemical parameters only. To test this hypothesis, the previously published method to predict lipid interactions from experimental f_u,mic_ data^12^ was implemented into the dynamic model (LDM-MB) and tested. The K_p_ prediction using the LDM-MB was improved in particular for gemfibrozil (**Table 2 & Figure 3**), but the model performance still did not meet acceptance criteria. The observed liver K_p_ remained significantly underpredicted (**Table 2)** with the predicted liver K_p_ values being on average 6.1-fold greater than the observed. The AFE and AAFE were 0.18 and 5.41 for the 4 drugs using the LDM-MB (**Table 2 & Figure 3**). Overall, none of the commonly considered distribution mechanisms and prediction methods captured the rat liver K_p_ values within 2-fold for the four drugs studied (**Figure 3**). This suggests that some distribution mechanisms in the liver are not captured in LDM-RR, LDM-AB, or LDM-MB.

### 3.2 Development of LDM-IPB and sensitivity analysis

We hypothesized that binding of the four test drugs (diclofenac, gemfibrozil, pioglitazone, and tolbutamide) to FABP1 may increase their liver partitioning and liver K_p_ values. To test this hypothesis, the LDM-MB was refined to incorporate human liver physiology and FABP1 expression in the human liver cytosol. The structure of the model is shown in **Figure 2**.

To examine the structural integrity of the model with and without intracellular protein binding, sensitivity analysis was conducted for human LDM-MB and LDM-IPB. The results of the sensitivity analysis of LDM-MB are shown in **Figure S2-1.** For the LDM-MB, logP and pKa, within the ranges considered, had minimal effect on liver K_p_ for the hypothetical weakly acidic drug. In contrast, as expected from pharmacokinetic principles and steady state assumptions, K_d,albumin_, f_u,p_, and f_u,mic_ had strong-to-moderate impact on liver K_p_ (**Figure S2-1**). For the LDM- IPB, the predicted liver K_p_ value was not sensitive to logP and pKa, but sensitive to f_u,p_, and f_u,mic_ similar to the LDM-MB results (**Figure 4**). In addition, intracellular FABP1 expression and K_d,FABP1_ had strong-to-moderate impact on liver K_p_ due to the effects on drug-FABP1 binding equilibrium.

**Figure 4.**
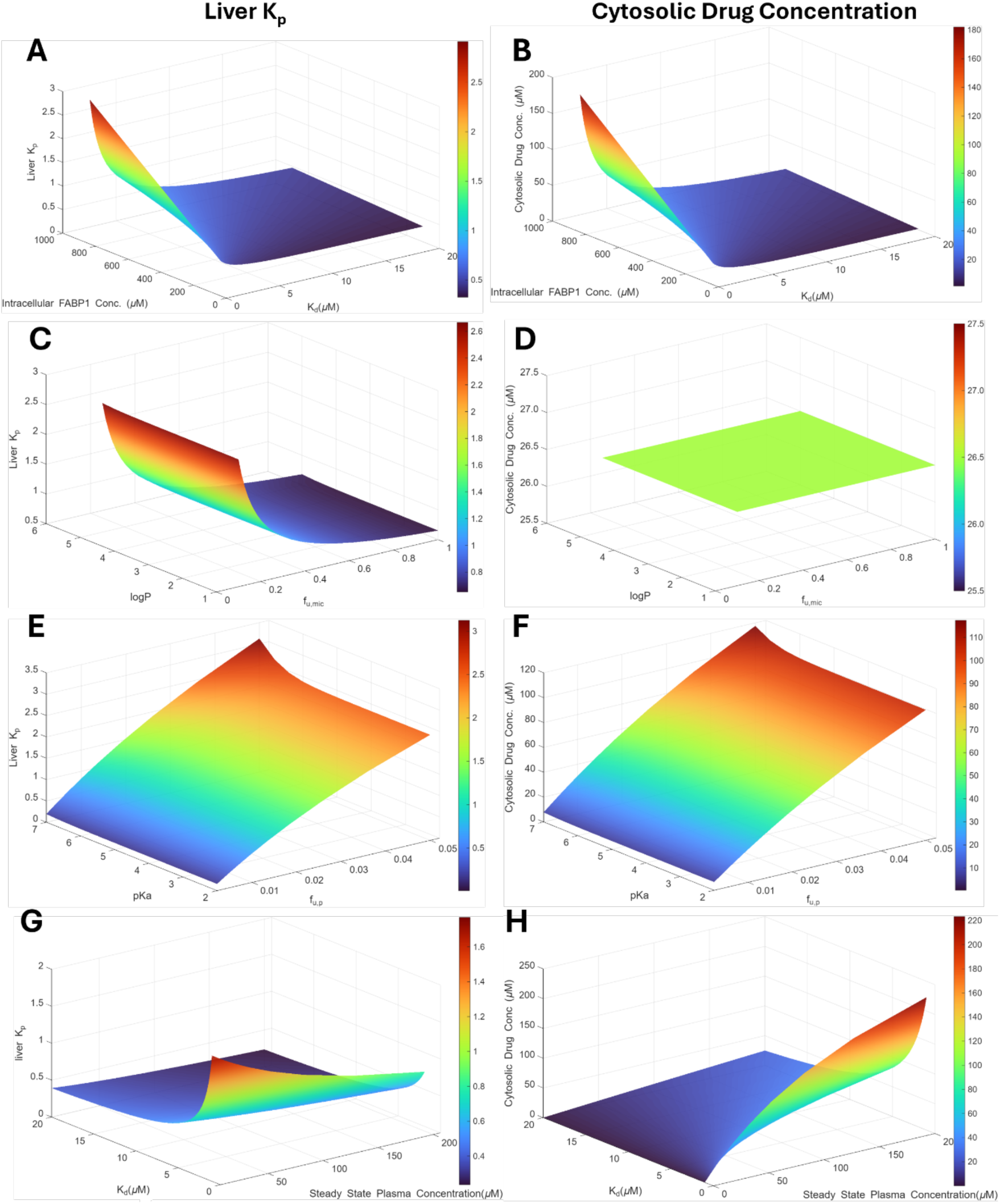
Sensitivity analysis of the developed human liver distribution model with intracellular protein binding (LDM-IPB). The sensitivity analysis was conducted using intracellular FABP1 expression of 399 µM and a hypothetical weakly acidic drug with fraction unbound in plasma (f_u,p_) of 0.01, logP of 3.84, blood-to-plasma ratio (BP) of 0.55, pKa of 8.14, drug-FABP1 equilibrium dissociation constant (K_d,FABP1_) of 3.6 µM, drug-albumin equilibrium dissociation constant (K_d,albumin_) of 2 µM with one binding site, and steady-state plasma concentration (C_ss_) of 40 µM. Panels A and B show the effects of intracellular FABP1 expression (100-900 µM) and K_d_ (1-20 µM) on liver K_p_ and cytosolic drug concentration. Panels C and D show the effects of logP (1-5) and f_u,mic_ (0.1-1) on liver K_p_ and cytosolic drug concentration. Panels E and F show the effects of pKa (2-7) and f_u,p_ (0.0001-0.05) on liver K_p_ and cytosolic drug concentration. Panels G and H show the effects of K_d_ (1-20 µM) and C_ss_ (0.1-200µM) on on liver K_p_ and cytosolic drug concentration.

In LDM-MB, C_cytosol_, the total cytosolic drug concentration, is the free drug concentration in plasma. Overall, C_cytosol_ was sensitive to f_u,p_ and C_ss_ while logP, f_u,mic_, and K_d,albumin_ did not impact free intracellular drug concentration as expected, demonstrating the robustness of the model (**Figure S2-1**). In LDM-IPB, C_cytosol_ was predicted as the sum of the free drug concentration and the concentration of drug-FABP1 complex. The C_cytosol_ was sensitive to FABP1 expression and K_d,FABP1_ in addition to being sensitive to f_u,p_ and C_ss_ similar to LDM-MB (**Figure 4**). Taken together, the sensitivity analyses demonstrate robustness of the LDM-IPB to predict liver distribution of drugs that bind to FABP1.

### 3.3 Measurement of FABP1 expression in 61 human livers

Simulation of the drug distribution into the liver in the presence of FABP1 relies on accurate quantification of FABP1 expression in the liver cytosol. Thus, the expression of FABP1 in the liver was quantified using targeted LC-MS/MS based peptide quantification approach (**Figure 5A&B**) and the measured expression level was scaled to the whole liver. The FABP1 expression in the human livers from 61 donors ranged between 121 and 436 nmol per gram of tissue. Considering liver f_ICW_ of 0.573^26^, the intracellular cytosolic concentrations of FABP1 in human livers were calculated to range from 211 µM to 760 µM with an average concentration of 399 µM, and a CV of 24.8% for all 61 human livers (**Figure 5C, Table S3-1**). The range between lowest and highest cytosolic FABP1 concentration quantified was about 3.6-fold. The quantified FABP1 concentrations were incorporated into the LDM-IPB to simulate interindividual variability in human liver K_p_ values.

**Figure 5.**
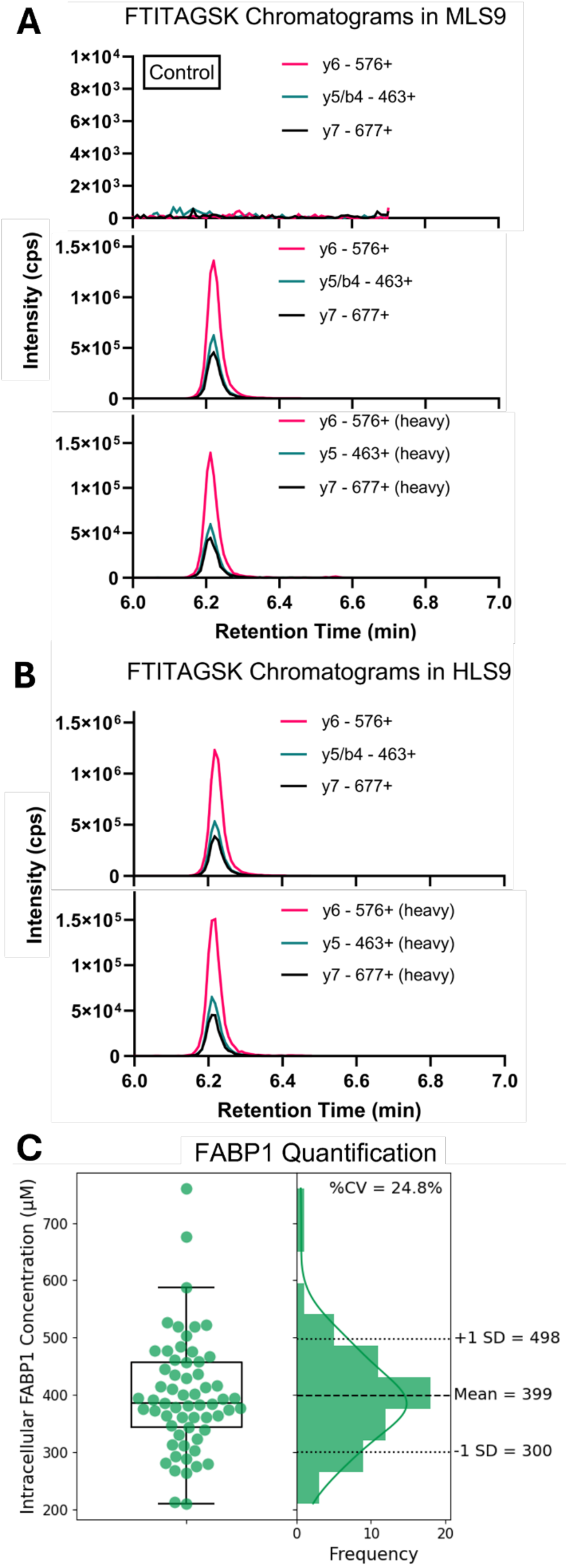
Representative chromatograms of the peptide FTITAGSK used for FABP1 quantification (A and B) and the distribution of FABP1 expression in 61 liver donors (C). Panel A shows the chromatograms of FTITAGSK and stable isotope–labeled FTITAGSK in MLS9 without (top) or with (middle and bottom) spiked human FABP1. Panel B shows the chromatograms of FTITAGSK and stable isotope–labeled FTITAGSK in pooled HLS9. Panel C shows the quantified FABP1 concentrations in intracellular water in individual donors.

### 3.4 LDM-IPB verification, comparison to other liver models and static predictions, and individual human liver K_p_ prediction

The human liver K_p_ was predicted for the 4 model drugs that bind to FABP1^15^ (diclofenac, gemfibrozil, pioglitazone, and tolbutamide) using the LDM-IPB. The predicted values were compared to K_p,scaled,human_ values (**Table 3**) and to the predicted K_p_ values using the static model or the dynamic LDMs adapted to human physiology (**Figure 6A-C**). The static equation and LDM-RR underpredicted the human liver K_p_ values for all four drugs with more than 10-fold average error. The liver K_p_ predictions were more accurate using LDM-AB and LDM-MB but still did not meet the 2-fold criteria. Using average FABP1 expression level of 399 µM in the model, the human liver K_p_ values predicted using the LDM-IPB passed the pre-determined two-fold model acceptance criterion for the 4 drugs (**Table 3**, **Figure 6C**). This shows a considerable improvement from the static prediction method and from the three LDMs (**Figure 6A-C**). Between the static methods and the LDM-IPB the AFE improved from 0.09 to 0.81 with average 8.5-fold improvement and the AAFE improved from 10.98 to 1.42 (**Table 3**, **Figure 6C**). The prediction accuracy improved by 4.3- to 13-fold when FABP1 binding was taken into consideration when compared to the static model that does not take drug-FABP1 binding into account (**Table 3**).

**Figure 6.**
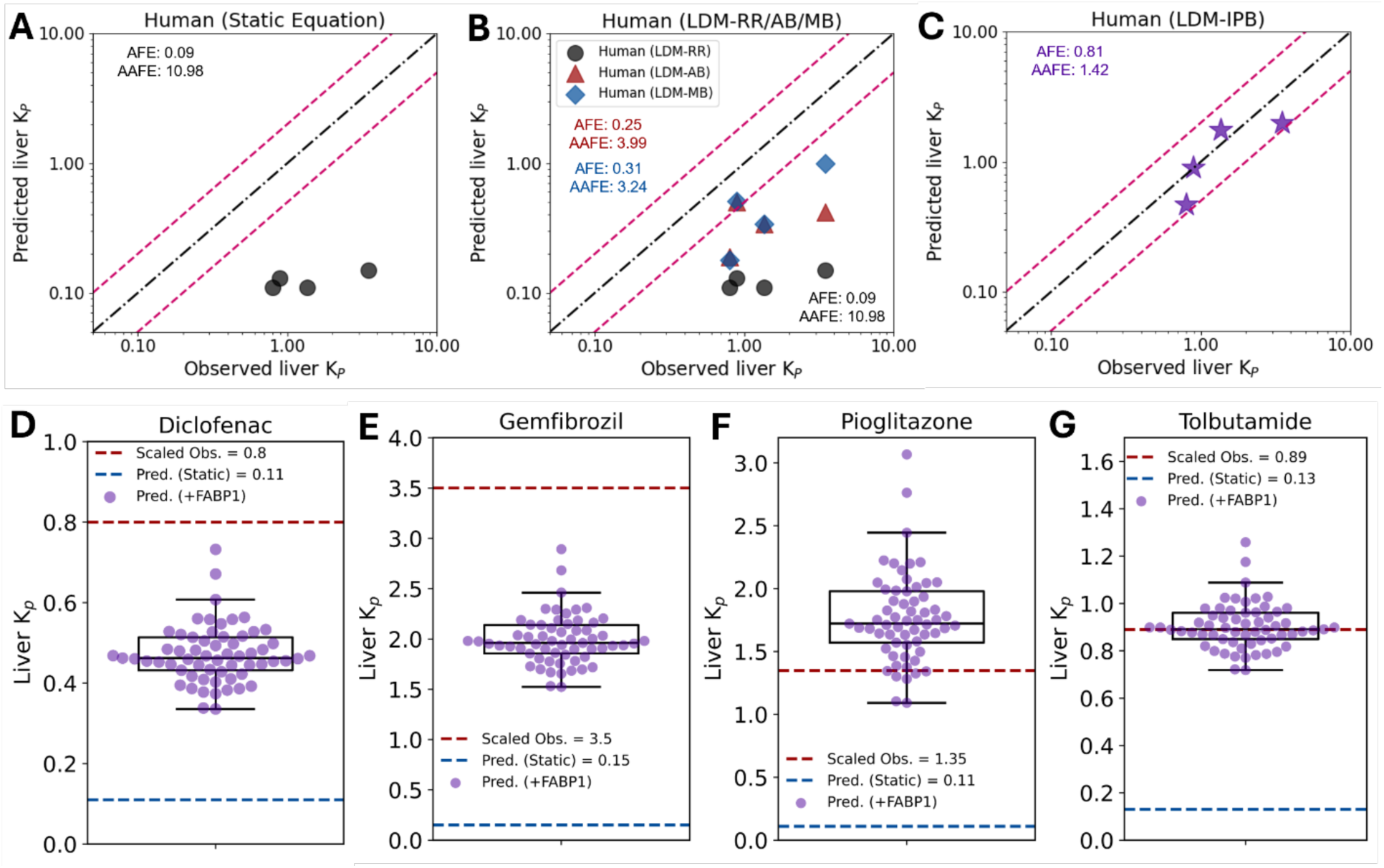
Simulation of liver K_p_ for diclofenac, gemfibrozil, pioglitazone, and tolbutamide. Panels A and B show the scaled human liver K_p_ from observed rat data versus predicted values using static equation or the LDM-RR/-AB/-MB without considering FABP1 binding. Panel C shows the scaled observed liver K_p_ in human versus simulated liver K_p_ after incorporating FABP1 binding with average FABP1 concentration of 399 µM using LDM-IPB. Panels D-G show box and whiskers plots of the simulated human liver K_p_ values for the four drugs in the 61 donors. The red dashed line represents the scaled observed liver K_p_ in human extrapolated from rat data. The blue dashed line indicates the predicted liver K_p_ using the static equation.

**Table 3.** Prediction of human liver K_p_ and cytosolic drug concentrations of the model compounds using the static method or LDM-IPB.

| Drug | Scaled Liver $K_p$ in Human | Predicted Liver $K_p$ | | Observed $C_{max}^{19}$ ($\mu M$) | Simulated $C_{ss}$ ($\mu M$) | Predicted $C_{cytosol}$ ($\mu M$) | |
| --- | --- | --- | --- | --- | --- | --- | --- |
|  |  | Static Equation<sup>3</sup> | LDM-IPB |  |  | Static Equation<sup>3</sup> | LDM-IPB |
| Diclofenac | 0.8 | 0.11 | 0.47 | 4.1 | 2.1 | 0.0065 | 1 |
| Gemfibrozil | 3.5 | 0.15 | 1.99 | 80 | 40 | 0.76 | 70 |
| Pioglitazone | 1.4 | 0.11 | 1.77 | 3.2 | 1.6 | 0.01 | 4 |
| Tolbutamide | 0.89 | 0.13 | 0.90 | 196 | 98 | 4 | 71 |
| AFE | - | 0.09 | 0.81 | - | - | - | - |
| AAFE | - | 10.98 | 1.42 | - | - | - | - |

Using the individual donor FABP1 expression level for the 61 donors while holding all the other model input parameters constant, the interindividual variability in liver K_p_ values for each of the four model drugs was simulated (**Figure 6D-G**). When the drug C_ss_ was set at 50% of the reported C_max_, 96.7%, 100%, 96.7%, and 100% of the individual simulated liver K_p_ values were within 2-fold of the K_p,scaled,human_ for diclofenac, gemfibrozil, pioglitazone, and tolbutamide, respectively (**Figure 6D-G**). The range between the minimum and maximum K_p_ values for a given drug was predicted to be 1.8- to 2.8-fold (11% to 20% CV) (**Figure 6D-G**).

### 3.5 Prediction of C_cytosol_ in human liver with and without considering drug-FABP1 binding

To evaluate the impact of drug-FABP1 binding on cytosolic drug concentrations, C_cytosol_ was simulated using the LDM-IPB and predicted using static equation (**Eq.28**) assuming no FABP1 in the ICW. Using the static equation under these assumptions, the predicted C_cytosol_ was equal to the unbound drug concentration in the cytosol. Due to high plasma protein binding for the four model drugs, C_cytosol_ values were low in comparison to total plasma concentrations (**Table 3**).

When average FABP1 expression level was incorporated into the model, population average C_cytosol_ (the sum of unbound drug concentration and drug-FABP1 complex concentration) was simulated. For the four model drugs, C_cytosol_ increased 18- to 400-fold (**Table 3**) when FABP1 binding was considered.

## 4. DISCUSSION

Prediction of tissue K_p_ values from *in vitro* data and physiochemical properties of the compound allows estimation of V_ss_ and tissue distribution kinetics without clinical data. When combined with clearance predictions, K_p_ predictions allow simulation of concentration-time profiles using PBPK models. Existing *in silico* prediction methods have good accuracy for K_p_ or K_p,u_ values for a subset of drugs. However, the K_p_ values for many drugs remain poorly predicted.^3,4,11,12,26^ This is a limitation during drug development as poor K_p_ predictions can lead to under or overprediction of estimated human half-life and clinical dose estimation. When detailed PBPK models are developed to estimate drug-drug interactions, disease impacts on drug disposition, or tissue/target site concentrations, poor predictions of K_p_ and V_ss_ values reduce the confidence in model performance and extrapolation. The data shown here suggest that incorporation of intracellular protein binding into PBPK models and tissue distribution predictions will significantly improve model quality and mechanistic underpinnings of simulation of distribution kinetics. Specifically, incorporation of FABP1 binding into tissue distribution was shown to allow accurate prediction of the extent of liver distribution of acidic drugs (**Figure 6C & Table 3**).

The developed LDM demonstrates the feasibility for a bottom-up dynamic prediction of tissue K_p_ for drugs using f_u,p_, logP, pKa, f_u,mic_, *in vitro* binding affinity, and protein expression data. The considerable error in predicting K_p_ values for weakly acidic drugs with underpredicted liver K_p_ was addressed by incorporating intracellular FABP1 binding into the dynamic models. While some prior studies showed that mechanisms such as plasma protein binding, ion partitioning, or lipid interaction can affect tissue distribution predictions,^3,4,11,12,26^ the current study provides insights for an additional intracellular protein binding concept to be integrated into the previously established framework. To address possible reasons for the underprediction in liver K_p_ observed, four different dynamic models were developed to simulate liver drug distribution. These models (**Figure 2**) demonstrate the possibility to integrate mechanistic distribution kinetics into a PBPK model instead of relying on static K_p_ values. LDM-RR as the dynamic version of Rodgers and Rowland method allows for prediction of tissue K_p_ for any small molecule of interest within a PBPK model. LDM-AB with the mechanistic albumin binding incorporated into the ECW demonstrated better prediction in drug-albumin partitioning than the traditional static model which back-calculates albumin binding affinity in the liver from f_u,p_. LDM-MB integrates drug-lipid partitioning using experimental liver microsomal binding data adapted from previously publihed methods^12^, allowing for easy prediction of drug distribution into tissue lipids. LDM-IPB provides new insights for understanding drug tissue distribution when intracellular protein binding is present *in vivo*. The dynamic models provide the first step in integrating mechanistic distribution kinetics into full-body PBPK models, allowing for more precise target site concentration prediction. With physiological parameter inputs for different tissues or species, the developed models can be adapted to other scenarios for distribution predictions, providing flexible applications.

We previously reported that diclofenac, gemfibrozil, pioglitazone, and tolbutamide form ternary complexes with human FABP1 *in vitro*, with K_d,FABP1_ values between 1 and 20 µM.^15^ The current model shows that despite these K_d,FABP1_ values being in the micromolar range, FABP1 binding can be substantial *in vivo* due to the high FABP1 intracellular concentration measured here for the human liver. This extent of binding is analogous to plasma albumin binding of the model drugs where the micromolar binding affinity with albumin translates to high protein binding in plasma or ECW. The substantial underprediction in liver K_p_ for drugs that bind to FABP1 and the significant improvement in prediction accuracy after taking FABP1 binding into account suggest that intracellular protein binding is important *in vivo* (**Table 2 & 3**).

Even with a modest binding affinity, drug-FABP1 binding can increase drug tissue concentration substantially, especially for weakly acidic drugs that typically have limited tissue distribution. The FABP1 binding in the cytosol can be pharmacologically and kinetically significant and affect predictions of clearance or pharmacodynamic outcomes. It has been suggested that FABP1 channels its ligands to enzymes and nuclear receptors.^15,17,41,42^ Therefore, if the drugs are bound to FABP1 in the cytosol this total rather than the unbound concentration is likely pharmacologically relevant. Accurate predictions of drug-FABP1 complex concentrations in the cytosol, thus, have downstream implications. Even though the free drug concentration is predicted to be the same in the presence and absence of FABP1, the overall C_cytosol_ can be orders of magnitudes higher if the drug binds to FABP1 in the liver than in the absence of FABP1 binding (**Table 3**). This is likely important for drugs such as pioglitazone and gemfibrozil that target peroxisome proliferator-activated receptors (PPARs) as FABP1 has been shown to channel its ligands to PPARs.^40^ Within this context, the total cytosolic concentration rather than unbound plasma concentration is likely pharmacodynamically relevant. For example, recently FABP1 binding was shown to stabilize acyl glucuronide from migration, potentially reducing the toxicological potential of the acyl glucuronide.^43^ Conversely, the high cytosolic concentrations may also be relevant for consideration of off-target toxicity via a variety of mechanisms.

Understanding the importance of FABP1 in modulating liver drug distribution requires knowledge of the expression of FABP1 in human liver. Past studies have estimated that FABP1 expression in the liver of preclinical species and in humans ranges from 0.1 mM to 1 mM.^17,44–48^ However, the methods used for quantification have been semiquantitative or qualitative, such as western blotting and immunohistochemistry, and the range of values is wide.^17,44–48^ This study is the first to quantify human FABP1 expression in the liver using targeted LC-MS/MS based proteomics approach with a reference protein standard included in the analysis. The measured population average (∼0.4 mM) of FABP1 expression is consistent with prior measurements of FABP1 cytosolic concentration. The 61 donors included in the FABP1 quantification had variety of liver pathologies including fatty liver, fibrosis, and acute injury potentially contributing to the population variability in FABP1 concentrations. The variability of FABP1 expressions within this liver bank is 3.6-fold. When compared to another study in which relative MS/MS was used to measure FABP1 expression in liver donors with liver disease, only up to 2-fold relative abundance difference was observed among liver donors with different disease progression.^44^ Due to the lack of calibration curves, absolute FABP1 expression level was not quantified. It is likely that the variability in FABP1 expression measured here translates to patient populations, and as such these data suggest that liver K_p_ will vary between individuals for drugs that bind to FABP1. Together with the knowledge of the interindividual variability in FABP1 expression, the developed LDM-IPB allows for simulation of the interindividual variability in liver K_p_ and in total C_cytosol_. Incorporating individual FABP1 expression levels into the LDM-IPB, population-based variability in drug distribution into the liver due to drug-FABP1 binding can also be simulated, allowing for prediction of interindividual variability in tissue distribution and V_ss_ (**Figure 6D-G**).

To our knowledge, this is the first study that examined the impact of intracellular protein binding on small molecule drug distribution integrating *in vitro* binding data, physiochemical properties of the drug, and protein quantification results into an *in silico* model. Many drugs bind to FABP1^13–15^, suggesting that the phenomenon described here is widespread. The LDM-IPB developed can be extrapolated to kidney and small intestine where considerable FABP1 expression is also seen in humans.^17^ In addition to FABP1, nine other FABPs are expressed in adipose, muscle, brain, heart, lung, and other organs.^17^ While the liver is a relatively small organ and hence typically does not have a major impact on overall total body V_ss_ prediction, tissues like adipose and muscle have relatively large volumes and K_p_ predictions for these organs impact overall V_ss_ predictions considerably. If FABP binding in these organs is similarly important as shown here for liver, intracellular protein binding will have a significant impact on overall V_ss_ prediction. Meanwhile, multiple FABPs can also be present in one tissue.^17^ The prevalence of FABPs in different organs in humans^17^ and the promiscuous and widespread binding of drugs to different isoforms of rat or human FABPs *in vitro*,^13–15^ suggest that the concept of intracellular protein binding is relevant to drug distribution to many organs. The developed LDM-IPB can be easily adapted to predict drug distribution in other tissues of interest to address this.

Mechanistically, the LDM-IPB can be integrated into a full-body PBPK model to simulate concentration-time profiles without using static tissue K_p_ values. This approach of modeling tissue distribution in dynamic fashion is important for drugs that bind tightly to albumin or intracellular binding proteins. Dynamic models allow simulation of circulating concentration dependent (saturable) distribution kinetics and nonlinear distribution phenomenon mechanistically. If a drug of interest binds to albumin or cytosolic proteins tightly *in vivo*, distribution kinetics is expected to be plasma concentration dependent. This is due to the equilibrium that must be restored among the free binding protein, the free drug, and the drug-binding protein complex formed in extracellular and intracellular space. The developed LDM-IPB allows simulation of such saturable phenomena for small molecule drugs.

In conclusion, this study shows that drugs that bind FABP1 *in vitro* will also bind FABP1 *in vivo* in the liver. The simulations show that FABP1 binding likely has a significant impact in the extent of distribution of drugs into the liver and suggest FABP isoforms in other tissues may further contribute to drug distribution kinetics. The developed dynamic models can be incorporated into whole body PBPK models and used to predict and study distribution kinetics and intracellular target binding.

## Data availability statement

The raw data that support the findings of this study are available from the corresponding author upon reasonable request. The model code is freely available on GitHub (https://github.com/Isoherranen-Lab/Liver-Distribution-Model).

## CRediT authorship contribution statement/Authorship contributions

**Yue Winnie Wen:** Writing – original draft, Visualization, Methodology, Investigation, Formal analysis, Funding acquisition, Conceptualization.

**Nina Isoherranen:** Writing – review & editing, Conceptualization, Supervision, Project administration, Methodology, Investigation, Funding acquisition.

## Acknowledgments

This work was supported in part by grants from the National Institutes of Health P01 DA032507, and R01 DK143511 and by the Drug Metabolism Transport and Systems Pharmacology Research Consortium (DMTSPR) at the University of Washington. NI is supported in part by the Milo Gibaldi Endowed Chair of Pharmaceutics to Department of Pharmaceutics, University of Washington.

## Nonstandard Abbreviations

FABP1, liver fatty acid binding protein; K_d_, equilibrium dissociation constant; K_p_, tissue-to-plasma partition coefficient; V_ss_, steady-state volume of distribution; PBPK, physiologically based pharmacokinetic model; AFE, average fold error; AAFE, absolute average fold error; LDM-RR, liver distribution model as a dynamic version of Rodgers and Rowland method; LDM-AB, liver distribution model with mechanistic albumin binding in extracellular water space; LDM-MB, liver distribution model with lipid partitioning predicted using microsomal binding data; LDM-IPB, liver distribution model with intracellular protein binding; LC-MS/MS, liquid chromatography-tandem mass spectrometry; f_u,p_, fraction unbound in plasma; f_u,mic_, fraction unbound in liver microsomes; K_d, albumin_, drug-albumin equilibrium dissociation constant; K_NL_, partition constant for neutral lipid binding; K_d, FABP1_, drug- FABP1 equilibrium dissociation constant; logP, log octanol-to-water partition coefficient; BP, blood-to-plasma ratio; C_max_, peak plasma concentrations; C_cytosol_, cytosolic drug concentration; QC, quality control; LLOQ, lower limit of quantification.

## Recommended section assignment

Drug Discovery and Translational Medicine or Metabolism, Transport, and Pharmacogenomics

