## Supplemental materials for "Development of Physiologically Based Liver Distribution Model that Incorporates Intracellular Lipid Partitioning and Binding to Fatty Acid Binding Protein 1 (FABP1)"

Figure S3-1. Detection of human specific peptides (A-D) of FABP1 from initial peptide screen following digestion of 200 nM FABP1 in 0.2 mg/mL mouse liver S9 (8 µg protein). 9

Figure S3-4. Trypsin digestion time course for potential surrogate peptides from 200 nM purified FABP1 digested in 4 µg mouse liver S9 fraction or 4 µg human liver S9 digested.12

### S1: Observed rat $K_p$ and drug specific parameters derived from literature.

**Table S1-1. Experimental information of liver distribution studies and liver $K_p$ values for diclofenac, gemfibrozil, pioglitazone, and tolbutamide in rats estimated from experimental data.**

| Drug | Dose and Route of Administration | Liver-to-Plasma Concentration Ratio (Time) | Estimated Liver $K_p$ |
|---|---|---|---|
| Atenolol[1] | 6 µmoles/kg every 60 min, five doses intraperitoneally, sixth dose intravenously | - | 2.21 (estimated from $AUC_{inf}$ ratio) |
| (R)-Propranolol[2] | 6.9 µg/min infusion for 4 and 8 h infusions to a total dose volume of 1 or 2 mL per animal | - | 5.56 (steady-state; corrected for hepatic extraction ratio) |
| (S)-Propranolol[2] | 6.9 µg/min infusion for 4 and 8 h infusions to a total dose volume of 1 or 2 mL per animal | - | 30 (steady-state; corrected for hepatic extraction ratio) |
| Diclofenac[3] | 40 mg/kg subcutaneous injection | - | 0.75 (estimated from $AUC_{0\text{-}inf}$ ratio) |
| Gemfibrozil[4] | $^{14}C$: 1 mg/kg IV injection | 3.2 (1 hr); 3 (4 hr); 3.5 (8 hr) | 3.2±0.3 |
| Pioglitazone[5] | $^{14}C$: 0.5 mg/kg oral | 1.21 (2 hr); 1.25 (6 hr); 1.48 (10 hr) | 1.3±0.1 |
| Tolbutamide[6] | A loading dose of 40.79 mg/kg followed by a constant rate infusion of 15.43 mg/kg per hour | - | 0.30±0.05 (steady-state) |

**Table S1-2. Comparison of hepatic extraction ratios for the model drugs in rats and humans.** Liver extraction ratio was calculated as $\frac{CL_{h,p}}{BP \cdot Q_h}$ where $CL_{h,p}$ is the hepatic clearance in plasma shown in the table below, BP is the blood-to-plasma ratio shown in Table 1, and $Q_h$ is hepatic blood flow (55 mL/min/kg[7] in rats and 21 mL/min/kg[7] in humans).

| | Rat hepatic clearance (mL/min/kg) | Rat liver extraction ratio[a] | Human hepatic clearance[b] (mL/min/kg) | Human liver extraction ratio[c] |
|---|---|---|---|---|
| Diclofenac | 14.4[8] | 0.48 | 4.2 ± 0.9[9] | 0.36 |
| Gemfibrozil | - | 0.09[10] | 1.7 ± 0.4[9] | 0.15 |
| Pioglitazone | 1[5] | 0.03 | 1.2 ± 1.7[9] | 0.10 |
| Tolbutamide | 0.23[11] | 0.007 | 0.22 ± 0.06[9] | 0.017 |

[a]Average 0.25-kg body weight was used for rat hepatic extraction ratio calculation except for gemfibrozil where extraction ratio was provided in the reference
[b]Hepatic clearance was assumed to be the same as total body clearance as renal clearance was negligible for all model drugs
[c]Average 70-kg body weight was used for human hepatic extraction ratio calculation

**Table S1-3. Physiological parameters used in the developed human and rat liver distribution models.** Physiological parameters are based on a 70-kilogram healthy adult human or 250-gram healthy adult rat.

| | Rat | Human |
|---|---|---|
| ***Liver Volume (L)***[12,13] | 0.00915 | 1.8 |
| Liver blood[12] | 0.00192 | 0.2 |
| Liver tissue[12] | 0.00723 | 1.6 |
| Liver – space of disse[12,14,15] | 0.00115 | 0.26 |
| Liver – intracellular space[12,14,15] | 0.00395 | 0.92 |
| ***Fraction of Liver Wet Weight*** | | |
| Water[14,15] | 0.705 | 0.73 |
| Neutral lipids ($f_{NL}$)[15] | 0.0138 | 0.0348 |
| Neutral phospholipids ($f_{NP}$)[15] | 0.0303 | 0.0252 |
| Extracellular water ($f_{ECW}$)[15] | 0.159 | 0.161 |
| Intracellular water ($f_{ICW}$)[15] | 0.546 | 0.573 |
| ***Fraction of Plasma Wet Weight*** | | |
| Neutral lipids ($f_{p,NL}$)[15] | 0.00149 | 0.0032 |
| Neutral phospholipids ($f_{p,NP}$)[15] | 0.00083 | 0.0021 |
| ***pH*** | | |
| Plasma[16] | 7.4 | 7.4 |
| Extracellular water[16] | 7.4 | 7.4 |
| Intracellular water[16] | 7.2 | 7.2 |
| ***Albumin Ratio*** | | |
| Extracellular water-to-plasma[17,18] | 0.54 | 0.63 |
| Liver-to-plasma[15,17] | 0.086 | 0.101[a] |
| ***Albumin Concentration (µM)*** | | |
| Plasma[19,20] | 409 | 640 |
| Liver – extracellular water[17–20] | 221 | 403[b] |

[a]Liver-to-plasma albumin ratio is calculated as $f_{ECW}$ × extracellular-to-plasma albumin ratio.
[b]Albumin concentration in the extracellular water space in liver is calculated as the plasma albumin concentration × extracellular water-to-plasma albumin ratio.

#### S2: Development of the liver distribution model (LDM) and sensitivity analysis

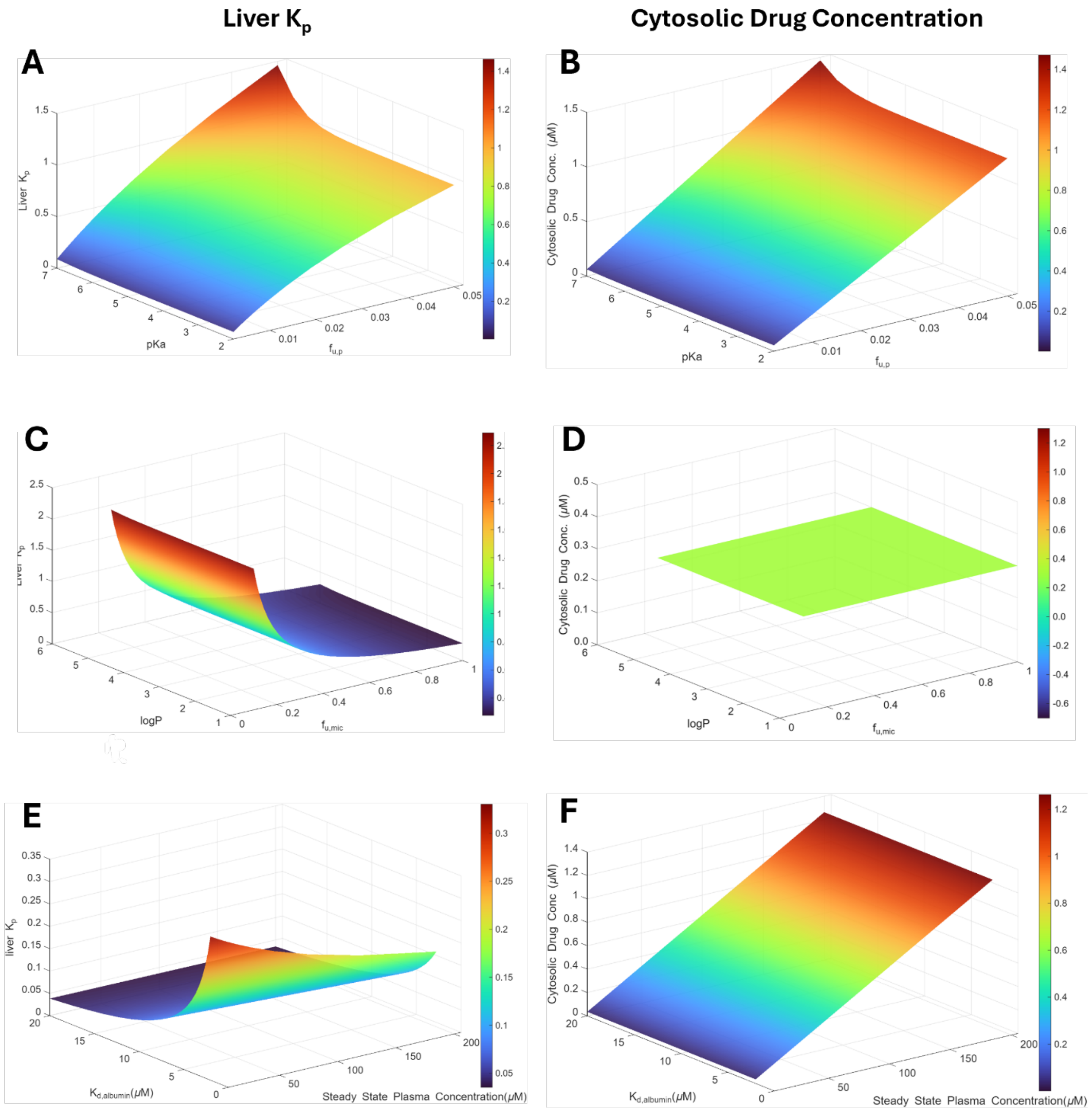

**Figure S2-1. Sensitivity analysis of the developed human LDM-MB.** The sensitivity analysis was conducted using a hypothetical weakly acidic drug with fraction unbound in plasma ($f_{u,p}$) of 0.01, logP of 3.84, blood-to-plasma ratio (B/P) of 0.55, pKa of 8.14, drug-albumin equilibrium dissociation constant ($K_{d,albumin}$) of 1 μM with one binding site, and steady-state plasma concentration ($C_{ss}$) of 40 μM. Panels A, C E show the effects of logP (1-5), $f_{u,mic}$ (0.1-1), pKa (2-7), $f_{u,p}$ (0.0001-0.05), $K_{d,albumin}$ (0.5-20 μM), and $C_{ss}$ (0.1-200μM) on liver $K_p$ prediction. Panel B, D, & F show the effects of the same sets of parameters on liver cytosolic drug concentration simulation.

### S3: Development and validation of liver FABP1 quantification method using surrogate peptides and LC-MS/MS

**Table S3-1. Liver weight processed, S9 yield, measured FABP1 concentration in S9, tissue FABP1 expression, and calculated human liver FABP1 intracellular concentrations in 61 liver donors extrapolated from fractional volume of intracellular water.**

| Donor ID | Tissue Weight (mg) | S9 Yield (mg S9/g liver) | FABP1 concentration in S9 (nmol FABP1/mg S9) | Tissue FABP1 Expression (nmol/g tissue) | Intracellular FABP1 Expression (µM) |
|---|---|---|---|---|---|
| HL-102 | 46.8 | 101 | 1.56 | 158 | 276 |
| HL-103 | 160.5 | 178 | 1.32 | 235 | 410 |
| HL-104 | 157.5 | 271 | 0.64 | 174 | 304 |
| HL-105 | 184.5 | 182 | 1.64 | 298 | 519 |
| HL-106 | 184.9 | 190 | 0.64 | 121 | 211 |
| HL-108 | 179.7 | 168 | 1.15 | 194 | 339 |
| HL-109 | 191.2 | 234 | 0.93 | 219 | 382 |
| HL-111 | 189.4 | 363 | 1.07 | 387 | 676 |
| HL-112 | 192.9 | 279 | 0.75 | 209 | 365 |
| HL-113 | 168.9 | 296 | 0.93 | 274 | 477 |
| HL-114 | 166.1 | 207 | 0.78 | 161 | 280 |
| HL-115 | 172.1 | 277 | 0.90 | 249 | 435 |
| HL-118 | 167.5 | 280 | 0.91 | 255 | 445 |
| HL-119 | 184.1 | 335 | 0.69 | 231 | 402 |
| HL-120 | 173.4 | 264 | 0.86 | 226 | 394 |
| HL-121 | 195.7 | 343 | 1.27 | 436 | 760 |
| HL-125 | 170.5 | 267 | 0.60 | 161 | 281 |
| HL-127 | 168.7 | 183 | 1.18 | 216 | 377 |
| HL-128 | 202.5 | 188 | 1.01 | 189 | 331 |
| HL-129 | 191.9 | 152 | 1.36 | 207 | 361 |
| HL-130 | 51.7 | 193 | 1.10 | 213 | 372 |
| HL-131 | 171.2 | 221 | 1.31 | 289 | 504 |
| HL-132 | 195.4 | 143 | 1.26 | 180 | 314 |
| HL-133 | 193.3 | 327 | 0.72 | 237 | 413 |
| HL-134 | 161.7 | 171 | 1.39 | 238 | 415 |
| HL-135 | 172.6 | 271 | 1.24 | 336 | 587 |
| HL-136 | 157.1 | 236 | 1.13 | 268 | 467 |
| HL-137 | 187.2 | 192 | 1.44 | 277 | 484 |
| HL-138 | 194.6 | 236 | 1.16 | 273 | 477 |
| HL-139 | 167.9 | 230 | 0.96 | 221 | 386 |
| HL-141 | 162.4 | 218 | 0.90 | 197 | 344 |
| HL-142 | 175.3 | 258 | 0.87 | 226 | 394 |
| HL-143 | 189.0 | 197 | 1.25 | 246 | 429 |
| HL-144 | 167.7 | 231 | 0.90 | 207 | 362 |
| HL-145 | 155.6 | 174 | 1.52 | 264 | 462 |
| HL-146 | 199.1 | 276 | 0.91 | 250 | 436 |
| HL-148 | 163.9 | 193 | 1.08 | 207 | 362 |
| HL-149 | 181.1 | 212 | 0.73 | 154 | 269 |
| HL-150 | 182.6 | 204 | 0.82 | 168 | 293 |
| HL-151 | 164.5 | 192 | 1.12 | 215 | 376 |
| HL-152 | 191.7 | 223 | 1.34 | 299 | 522 |
| HL-153 | 178.7 | 235 | 0.94 | 220 | 384 |
| HL-154 | 188.1 | 210 | 1.00 | 209 | 364 |
| HL-155 | 151.8 | 229 | 1.04 | 239 | 417 |
| HL-156 | 174.6 | 209 | 1.03 | 215 | 375 |
| HL-157 | 182.2 | 194 | 0.96 | 185 | 323 |
| HL-158 | 191.9 | 304 | 0.86 | 262 | 458 |
| HL-159 | 198.5 | 202 | 1.09 | 220 | 383 |
| HL-160 | 197.2 | 248 | 0.91 | 226 | 394 |
| HL-161 | 197.2 | 291 | 1.04 | 301 | 526 |
| HL-162 | 180.3 | 243 | 0.94 | 229 | 400 |
| HL-163 | 192.7 | 344 | 0.76 | 263 | 459 |
| HL-164 | 159.0 | 240 | 0.69 | 166 | 289 |
| HL-165 | 177.5 | 216 | 1.00 | 217 | 378 |
| HL-166 | 190.3 | 229 | 1.30 | 297 | 519 |
| HL-167 | 161.8 | 193 | 0.78 | 151 | 263 |
| HL-168 | 164.0 | 213 | 0.93 | 199 | 346 |
| HL-169 | 184.1 | 171 | 1.05 | 179 | 312 |
| HL-170 | 169.7 | 140 | 0.88 | 123 | 214 |
| HL-171 | 171.5 | 208 | 1.08 | 225 | 392 |
| HL-172 | 179.6 | 246 | 1.11 | 273 | 476 |

##### *Method S3-1: Expression and purification of human FABP1*

Hexa-histidine-tagged (his-tag) human FABP1 was expressed in Rosetta 2 *E. coli* (Novagen, Madison, WI) and purified as previously described with minor modifications.[21] Briefly, his-tagged FABP1 was purified using a 1 mL HisTrap HP affinity column (GE Healthcare, Chicago, IL). The his-tag was cleaved using thrombin (0.01 U thrombin per 10 mg FABP1) after 16-hr incubation on ice. Cleaved FABP1 was buffer exchanged using a Superdex 75 (Marlborough, MA) column equilibrated with 50 mM potassium phosphate, 100 mM KCl pH 7.4. Delipidation of the cleaved FABP1 was performed using three rounds of butanol treatment (5 mL butanol to 5 mL cleaved FABP1) and 30-minute incubation with preconditioned Lipidex-5000 (Perkin Elmer Inc., Waltham, MA, USA; 0.1g per 1 mg of FBAP1) in 50 mM potassium phosphate, 100 mM KCl pH 7.4 as previously described[21]. The concentration of purified FABP1 was quantified via bicinchoninic acid (BCA) assay (Pierce, Waltham, MA). The delipidated FABP1 was stored on ice in 50 mM potassium phosphate, 100 mM KCl, pH 7.4 at 4°C or flash frozen with addition of 5% glycerol and stored at -80°C for protein quantification experiments.

##### *Method S3-2: Human liver donors and preparation of S9 fractions*

Human liver samples from individual donors (n=61) were obtained from the University of Washington (UW), human liver bank. Donors' ages ranged from 7 to 68 years. Donors included 28 females and 33 males, with 57 Caucasian, 1 Asian, and 3 Black or African American. According to surgeon's notes at organ collection, 27 livers had normal pathology, with the rest presenting various pathologies including fatty, acute injury, fibrotic, iron deposition or moderate microvesicular fat.

Human liver S9 fractions (HLS9) and mouse liver S9 fractions (MLS9) were prepared using standard methods. Briefly, liver tissue was cut and the aliquot weighted (46.8–202.5 mg) and transferred into a tissue homogenizing tube with ceramic beads. A constant ratio (5-fold v/w) of homogenizing buffer (50 mM Kpi, 250 mM sucrose, 50 mM KCl, pH 7.4 with EDTA-free protease inhibitor cocktail tablet, Millipore-Sigma, St. Louis, MO) to liver tissue was added directly to the tissue homogenizing tubes. The tissue was then homogenized using Omni Bead Ruptor 24 (Kennesaw, GA) using five 20-second disruption cycles at 5.00 m/s with a 3-second rest between cycles between -15 ºC to -10 ºC. The homogenates were transferred to new 1.7 mL protein low-binding microcentrifuge tubes and centrifuged at 9,000 × g for 30 minutes at 4 ºC to remove unbroken cells, debris and large organelles. Avoiding the top fat layer if present, the middle supernatant containing cytosol and microsomes was transferred to a new 1.7 mL low-binding microcentrifuge tube as the S9 fraction. Protein concentrations were measured using BCA assay (Thermo Fisher, Waltham, MA) and the S9 fractions aliquoted and stored at -80 ºC .

##### *Method S3-3: Selection of surrogate peptides and UHPLC-MS/MS method*

The amino acid sequence for human FABP1 in FASTA format was retrieved from UniProtKB database (P07148 · FABPL_HUMAN). To screen for potential tryptic peptides of FABP1, FABP1 was digested *in silico* with trypsin using Skyline-daily (25.1.1.206). Human and mouse proteomes were acquired from UniProtKB database in FASTA format and used as background proteome. Only peptides that are unique to FABP1 and specific to human were included in downstream analysis. Peptides with 5–25 amino acids and predicted *m/z* range of 50–1200 were considered. Peptides containing cysteine or methionine residues were excluded due to potential modification. Precursor ions with 2 or 3 charges, fragmentation (b and y ions with 1 or 2 charges), declustering potential and collision energies were predicted in Skyline-daily for transition screening and method development.

To test for detection, fragmentation, and analytical sensitivity of the candidate peptides, purified FABP1 (200 nM) was digested in 100 mM ammonium bicarbonate (pH 7.8) or 0.2 mg/mL MLS9 as previously described for 20-hr[22]. Detected unique and human specific peptides with top three transitions ranked by peak intensity for each peptide were included in the method for further analysis (**Figure S3-1, Table S3-2, & Figure S3-2**). Peptide with the highest summed peak area of the three transitions and no interference with the MLS9 blank was selected as the quantitative peptide for FABP1 quantification.

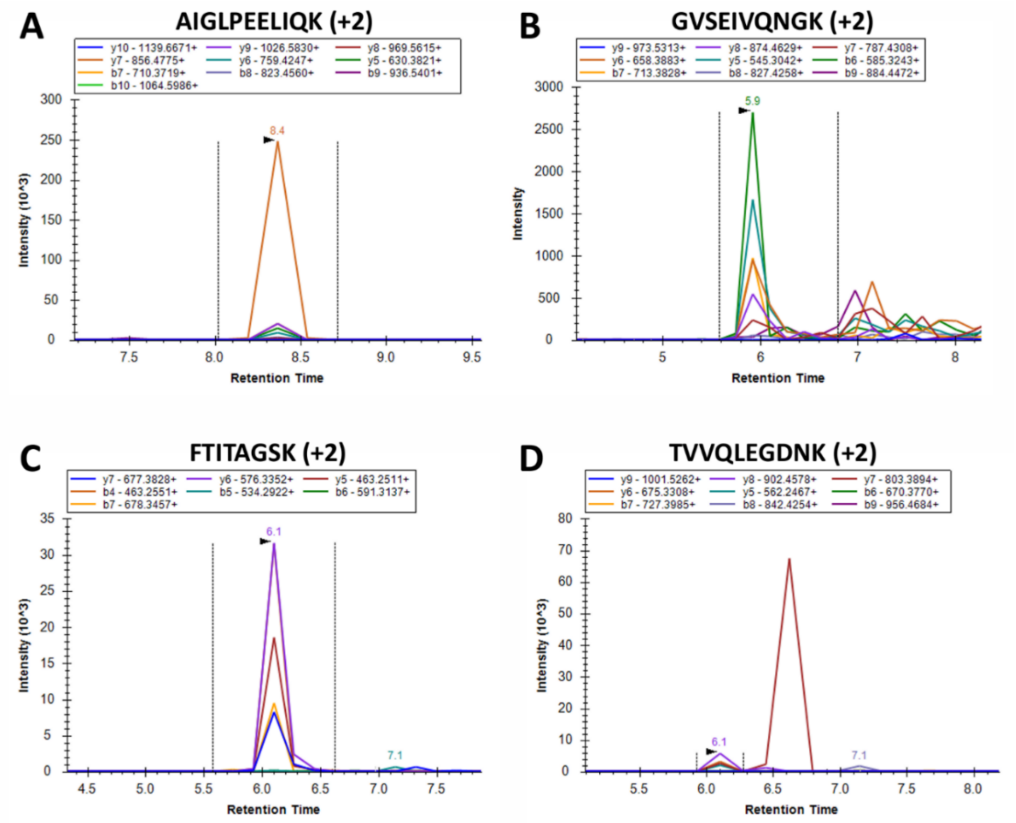

**Figure S3-1. Detection of human specific peptides (A-D) of FABP1 from initial peptide screen following digestion of 200 nM FABP1 in 0.2 mg/mL mouse liver S9 (8 µg protein).** The legends list the specific b or y ion, the charge state, and the MS/MS transitions monitored. Figure created with Skyline.

**Table S3-2. Peptide sequences and MS parameters of unique and human specific peptide selected for FABP1 quantification or monitored using LC-MS/MS.**

| Protein | Peptide | Precursor Ion (*m/z*) | Fragment Ion (*m/z*) | DP (V) | CE (V) |
|---|---|---|---|---|---|
| FABP1 | FTITAGSK (+2) | 413 | $y6^+$ 576 | 95 | 16 |
| | | | $y7^+$ 678 | 95 | 22 |
| | | | $y5^+/b4^+$ 463 | 95 | 22 |
| | F[$^{13}C_9{}^{15}N$]TITAGSK (+2) | 417 | $y6^+$ 576 | 95 | 16 |
| | | | $y5^+/b4^+$ 463 | 95 | 22 |
| | | | $y7^+$ 677 | 95 | 22 |
| **Monitored but not used for quantification** | | | | | |
| FABP1 | AIGLPEELIQK (+2) | 606 | $y7^+$ 856 | 80 | 29 |
| | | | $y5^+$ 630 | 80 | 29 |
| | | | $b7^+$ 710 | 80 | 29 |
| | GVSEIVQNGK (+2) | 516 | $b6^+$ 585 | 80 | 24 |
| | | | $b7^+$ 713 | 80 | 24 |
| | | | $y5^+$ 545 | 80 | 24 |
| | TVVQLEGDNK (+2) | 552 | $y8^+$ 903 | 80 | 26 |
| | | | $y6^+$ 675 | 80 | 26 |
| | | | $y5^+$ 562 | 80 | 26 |

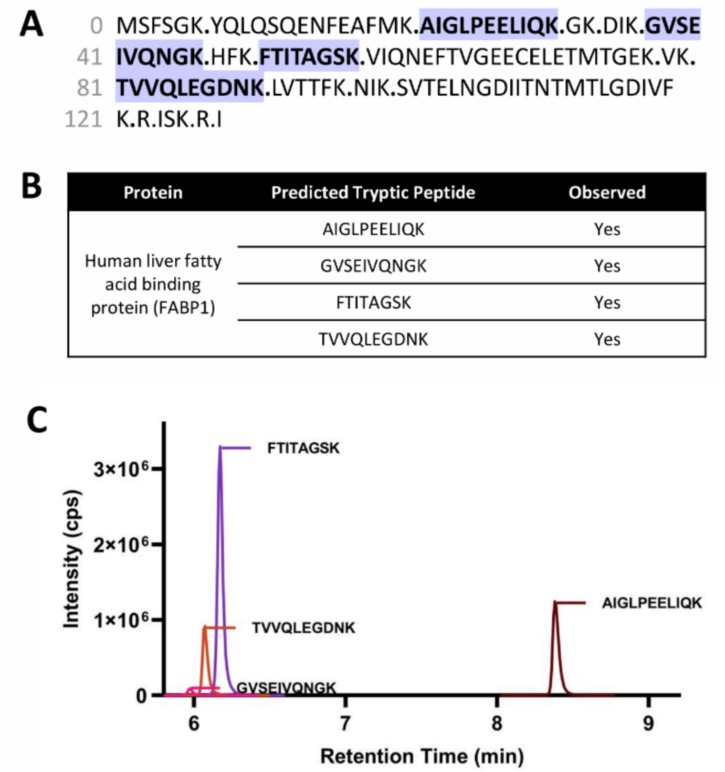

**Figure S3-2. Identification and selection of FABP1 surrogate peptides.** (A&B) Amino acid sequence of FABP1 acquired from UniProt.com. The trypsin cut site was shown with dots. The human specific and unique tryptic peptides were highlighted in purple. (C) Representative summed chromatograms of observed peptides following 20-hr digestion of 200 nM purified FABP1 in ammonium bicarbonate (pH 7.8).

##### *Method S3-4: Selection of digestion condition, digestion time, and evaluation of matrix effect*

The choice of detergent, reducing agent, and when to add stable isotope labeled internal standard peptide on digestion was assessed. Surfactant and denaturation effect on digestion efficiency was tested by digesting purified FABP1 (200 nM) in 0.2 mg/mL MLS9 and 0.2 mg/mL HLS9 in parallel with either 10 µL of 100 mM ammonium bicarbonate buffer (pH 7.8) or 10% (w/v) sodium deoxycholate in 100 mM ammonium bicarbonate buffer (pH 7.8). The experiments were conducted in six replicates in parallel followed by 95 ºC incubation for 5 minutes. The trypsin digestion time course experiment was conducted at 1:10 trypsin/protein (w/w) ratio for 20 hours. The reduction, alkylation, and quenching the digestion with acetonitrile containing 8% TFA was completed as described in the methods section. The peak area of the summed MS/MS transitions for the surrogate peptide FTITAGSK was compared between conditions (**Figure S3-3A**).

To evaluate the effect of time to add heavy labeled peptide F[$^{13}C_9{}^{15}N$]TITAGSK, 4 µL of 2,000 nM F[$^{13}C_9{}^{15}N$]TITAGSK diluted in 100 mM ammonium bicarbonate pH 7.8, was added to 0.2 mg/mL MLS9 with 200 nM FABP1 or to 0.2 mg/mL HLS9 at the beginning of the digestion in triplicates. The digestion was quenched with acetonitrile containing 8% TFA after 20 hours. In parallel, 4 µL of 100 mM ammonium bicarbonate (pH 7.8) was added instead at the beginning of the digestion in triplicates and the digestion was quenched with acetonitrile containing 8% TFA and 200 nM F[$^{13}C_9{}^{15}N$]TITAGSK with the remaining steps unchanged. The peak areas of the summed MS/MS transitions for F[$^{13}C_9{}^{15}N$]TITAGSK were compared between groups (**Figure S3-3B**).

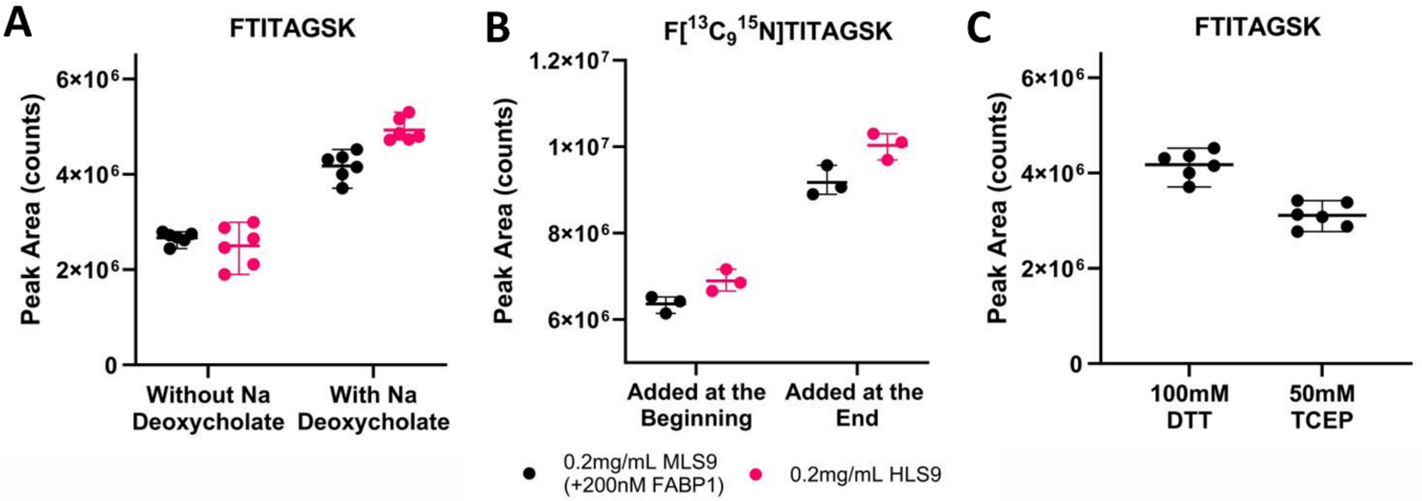

**Figure S3-3. Optimization of experimental conditions for FABP1 digestion.** (A) Peak area sum of multiple transitions monitored for FTITAGSK peptide as listed in Table S3-1. Samples were digested in parallel with either 10 µL of 10% sodium deoxycholate dissolved in 100 mM ammonium bicarbonate (pH 7.8) or 100 mM ammonium bicarbonate alone (pH 7.8). (B) Peak area sum of multiple transitions monitored for F[$^{13}C_9{}^{15}N$]TITAGSK peptide as listed in Table S3-1. 200 nM F[$^{13}C_9{}^{15}N$]TITAGSK was either added at the beginning of the digestion or at the end while keeping the rest of the protocol unchanged. (C) Peak area sum of multiple transitions monitored for FTITAGSK peptide as listed in Table S3-1. Samples were digested in parallel with either 8 µL of fresh 100 mM DTT or 50 mM TCEP dissolved in 100 mM ammonium bicarbonate (pH 7.8) added to 40 µL samples.

The effect of reduction with dithiothreitol (DTT) or tris(2-carboxyethyl)phosphine (TCEP) was evaluated by digesting 0.2 mg/mL HLS9 in parallel with either 8 µL of fresh 100 mM DTT or 50 mM TCEP dissolved in 100 mM ammonium bicarbonate (pH 7.8). The rest of the protocol was kept the same between digestions. The peak areas of the summed MS/MS transitions for FTITAGSK were used for reducing agent effect comparison (**Figure S3-3C**).

To select for the best digestion time, 0.2 mg/mL HLS9 and purified FABP1 (200 nM) spiked into a solution containing 0.2 mg/mL MLS9 both containing 30 nM F[$^{13}C_9{}^{15}N$]TITAGSK internal standards were digested in parallel in triplicates for 0.5-, 2-, 8-, 16-, and 24-hour (**Figure S3-4A-B, & D-F**). The rest of the protocol was kept unchanged as described in the method section. Summed peak area was acquired for each peptide for evaluation of digestion efficiency and completeness. To further optimize the digestion time, a subsequent interpolation digestion was performed for 2, 5 and 8 hours to find the optimum digestion time (**Figure S3-4C**).

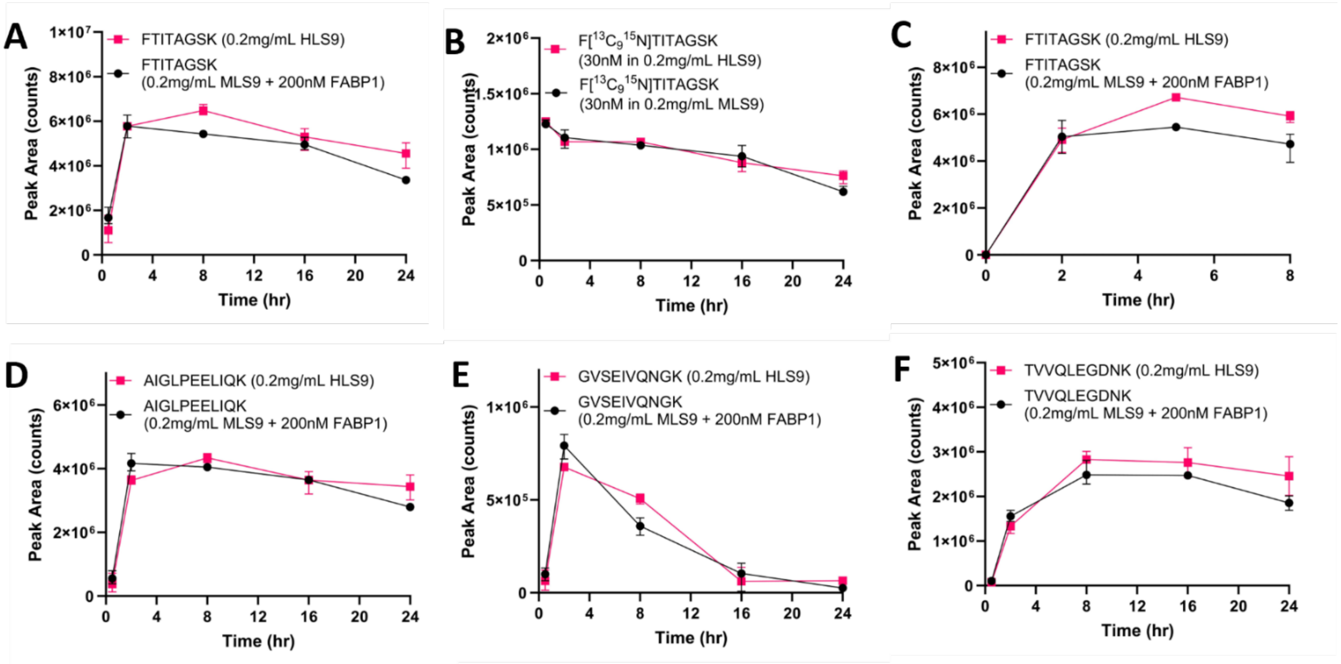

**Figure S3-4. Trypsin digestion time course for potential surrogate peptides from 200 nM purified FABP1 digested in 4 µg mouse liver S9 fraction or 4 µg human liver S9 digested.** 30 nM F[$^{13}C_9{}^{15}N$]TITAGSK was added at the beginning of the digestion in each sample digested. Panels A & B-F showed peak area sum of multiple transitions monitored for each peptide as listed in Table S3-1 at 0.5-, 2-, 8-, 16-, and 24-hour. Panel C showed peak area sum of multiple transitions monitored for FTITAGSK as listed in Table S3-1 at 2-, 5-, and 8-hour for determination of the best digestion time for FABP1 quantification.

To evaluate matrix effects, 0.2 mg/mL HLS9 pool and 0.2 mg/mL MLS9 matrix were digested for 5 hours. The digested HLS9 samples were serially diluted into digested MLS9 samples or 100 mM ammonium bicarbonate (pH 7.8) ranging from 0% to 100% HLS9. The linearity of the summed peptide peak area against dilution ratio was assessed using a simple linear regression (**Figure S3-5**).

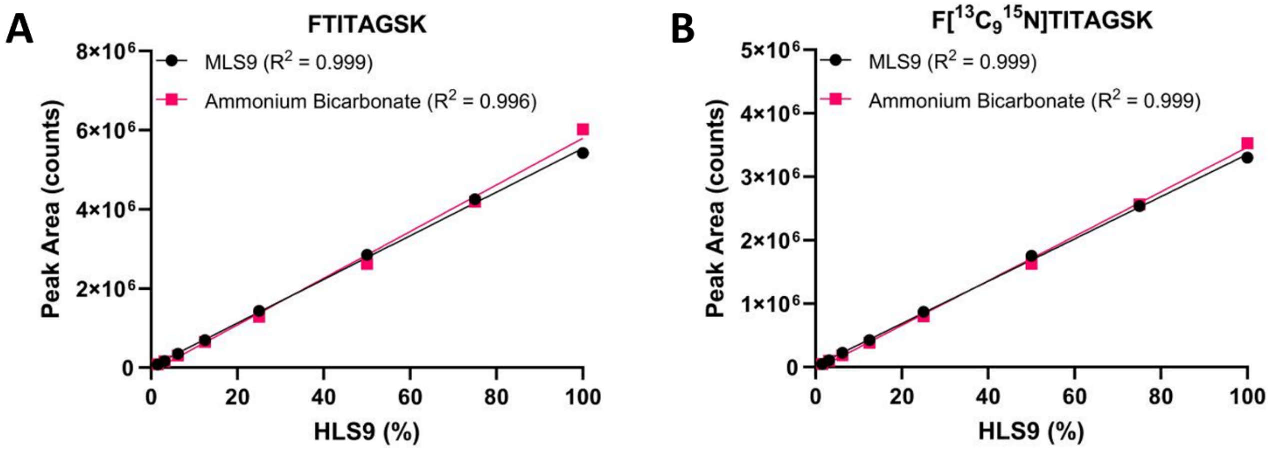

**Figure S3-5. Linearity of FTITAGSK and F[13C915N]TITAGSK peptide signal response on mass spectrometer.** Digested 0.2 mg/mL human liver S9 fraction (4 µg protein) was diluted to 0.2 mg/mL mouse liver S9 fraction (4 µg protein) digested or ammonium bicarbonate (pH 7.8). The $R^2$ values shown were from an unweighted linear regression.

**Table S3-3. Method validation data.** Accuracy (%error) and precision (% CV) of FABP1 quantification using FTITAGSK as a surrogate standard for quality control (QC) samples across 10 separate days. The % error was calculated as observed /nominal concentration for n=5 at each QC level for 4 separate runs. Intra-day variance was calculated from 5 replicates within a day and the average intra-day % CV for each QC level is reported. Stability of the surrogate peptide was determined following two freeze-thaw cycles at -20°C and the average from the 2 cycles is reported. Stability of the surrogate peptide in the autosampler was determined following 24-hr and 40-hr at 10°C and the average from the 2 runs is reported. Pooled QC was prepared as a mixture of HLS9 made from 61 donors. LLOQ: lower limit of quantification; LQC: low QC; MQC: middle QC; HQC: high QC.

| QC | Nominal Conc. (nM) | Inter-day error (%) | Inter-day variance (% CV) | Intra-day % CV | Freeze-thaw error (%) | Autosampler error (%) |
|---|---|---|---|---|---|---|
| LLOQ | 50 | 11 | 2.7 | 1.6 | 11 | 15 |
| LQC | 120 | -8.0 | 3.4 | 3.1 | -8.9 | -8.9 |
| MQC | 320 | -5.3 | 6.8 | 6.0 | -11 | -5.5 |
| HQC | 540 | 4.1 | 10 | 7.9 | 4.0 | 9.2 |
| Pooled QC | - | - | 13 | 5.9 | - | - |
